# Ribosome-Associated GTPase Drg1 Links Cytosolic Translation to Mitochondrial Homeostasis

**DOI:** 10.64898/2026.02.23.707305

**Authors:** Sounak Saha, Aryaman Bhattacharya, Hong Jin

**Affiliations:** Department of Biochemistry, University of Illinois at Urbana-Champaign, 600 S. Mathews Avenue, Urbana, IL 61801; Center for Biophysics and Quantitative Biology, University of Illinois at Urbana-Champaign, 600 S. Mathews Avenue, Urbana, IL 61801; Carl R. Woese Institute for Genomic Biology, 1206 West Gregory Drive, University of Illinois at Urbana-Champaign, Urbana, IL 61801

**Keywords:** Ribosome, Translation, GTPase, Developmentally-regulated GTPase, Drg, Mitochondria, Protein Homeostasis

## Abstract

Coordination of the functionalities among key organelles is essential for maintaining the complexity and adaptability of the eukaryotic cell. Functionally versatile proteins, such as GTPases, often assume important roles in this process. A ribosome-binding GTPase called the developmentally regulated GTP-binding (Drg) protein is a conserved family of GTPases that promotes protein synthesis and regulates cytoskeletal dynamics, suggesting their importance for cell growth and proliferation. Mitochondrial proteostasis requires coordination between cytosolic translation and the targeting, import, and assembly of nuclear-encoded mitochondrial proteins. However, the ribosome-associated factors that support this coordination remain incompletely understood. Here, we identify the conserved GTPase Drg1, one of the two Drg paralogues in eukaryotic cells, as a translation factor required for mitochondrial homeostasis and cellular fitness. Human Drg1 and its cofactor Dfrp1 are associated with translating ribosomes and co-sedimented with polysomes. Deletion of Drg1 in HEK293T cells caused altered mitochondrial architecture, reduced mitochondrial membrane potential and ATP production, and impaired expression of a mitochondrially targeted reporter. Re-expression of wild-type Drg1 restored ATP production and reporter expression, whereas a GTPase-deficient Drg1 mutant failed to provide comparable rescue in ATP production, demonstrating that Drg1’s catalytic activity is required for its cellular function. Mitochondria-directed APEX2 proximity proteomics further connected Drg1-dependent translation to proteins involved in mitochondrial protein import, respiratory-chain function, and mitochondrial protein synthesis. Reporter analyses indicated that Drg1 loss preferentially compromises productive translation toward the 3′ region of the coding sequence. Together, these findings establish Drg1 as a ribosome-associated translation factor that supports the synthesis and homeostasis of proteins required for mitochondrial function. At the molecular level, loss of cellular homeostasis that is caused by compromised translation of cytoplasmically synthesized proteins functionally related to mitochondria underpins the observed phenomenon. Our results reveal a previously underappreciated connection between the cytosolic translation machinery and mitochondrial maintenance and provide a framework for understanding how defects in a conserved translational GTPase impair cellular energy metabolism and physiology.

## Introduction

Proteins are fundamental macromolecules that carry out virtually all essential cellular functions. Protein synthesis—the process of generating proteins from the genetic information encoded in nucleic acids—is therefore indispensable for life. This process occurs on the ribosome, a remarkable molecular machine, and is tightly coordinated with the physiological, metabolic, and energetic states of the cell (Biffo et al., 2024; Dazert and Hall, 2011). How cells produce, target, and degrade proteins to maintain homeostasis and to ensure proper cellular function is a fundamental question in molecular biology. Eukaryotic organelles, such as the nucleus, endoplasmic reticulum, and mitochondrion, each possess distinct proteomes but have either a limited or complete lack of protein synthesis capacity. As a result, nascent proteins essential for organelle function must be synthesized in the cytoplasm and subsequently delivered to their target organelles. This process can occur through three general mechanisms: mRNA localization followed by local translation (Fazal et al., 2019; Gonzalez et al., 1999; Kislauskis et al., 1994; Padron et al., 2019; Ross et al., 1997), co-translational targeting (Fazal et al., 2019; Gold et al., 2017; Jan et al., 2014; Luo et al., 2025; Zhu et al., 2025), or post-translational targeting (Chacinska et al., 2009; Neupert and Herrmann, 2007). The translation machinery temporarily pauses during folding or co-translational targeting, a process particularly important to import large proteins into organelles (Pechmann et al., 2014; Pechmann and Frydman, 2013; Stein and Frydman, 2019; Yu et al., 2015). Conversely, to prevent cytotoxicity, mistargeted proteins and stalled translation machinery must be recognized and eliminated through quality control pathways (Brandman and Hegde, 2016; Doma and Parker, 2006; Inada and Beckmann, 2024; Joazeiro, 2019; Sitron and Brandman, 2020). This creates a fundamental challenge: how can the cell distinguish functional ribosomal pauses, which naturally occur during co-translational processes such as protein folding and membrane translocation, from pathological stalls that signal defective translation?

Recently, we have reported that a highly conserved family of GTPases called ribosome-binding GTPase (Rbg) in yeast or developmentally regulated GTP-binding proteins (Drg) in humans is critical for translation (Zeng et al., 2021). Rbg/Drg GTPase plays a critical role in promoting protein synthesis in paused ribosomes and protecting protein synthesis by preventing inappropriate activation of quality control pathways (Saha et al., 2026). Rbg/Drg proteins are an ancient family of essential GTPases that are highly expressed in actively growing and developing cells across plants, animals, and humans. These proteins were initially named “*developmentally regulated*” upon their discovery due to their distinctive expression pattern in neuronal precursor cells in the developing mouse brain (Kumar et al., 1992). However, our recent phylogenetic analyses reveal that Obg/Rbg/Drg GTPases are conserved across all three domains of life (Saha et al., 2026). All known eukaryotes encode two paralogs, Drg1 and Drg2, whereas archaea contain only a single Drg protein (Li and Trueb, 2000), which likely represents the ancestral form of its eukaryotic counterparts. Structurally, Drg proteins share a highly conserved G domain composed of five canonical G motifs that enable GTP binding and hydrolysis. Additionally, modern Drg proteins possess three distinct domains: an N-terminal helix-turn-helix (HTH) domain, a central S5D2L (S5 Domain 2-like) domain inserted between the G motifs, and a C-terminal TGS domain (stands for *Threonyl-tRNA synthetase ThrRS*, GTPases, and *SpoT*) (**Supplemental Figure S1**). We demonstrated that Drg functions as a translation factor that binds to translating ribosomes and promotes protein synthesis when ribosomes experience translational slowing on mRNA (Zeng et al., 2021). This role is evolutionarily conserved (Saha et al., 2026), suggesting Drg as a critical regulator of translation dynamics across diverse organisms. Consistent with an essential physiological function, biallelic loss-of-function variants in human Drg1 cause a severe neurodevelopmental disorder characterized by developmental delay, microcephaly, short stature, and craniofacial abnormalities, while targeted disruption of Drg1 in mice causes preweaning lethality (Westrip et al., 2023). Nevertheless, the cellular pathways that are particularly dependent on mammalian Drg1 activity remain poorly defined.

Data from our earlier genomic study revealed that proteins functionally associated with mitochondria were among the most significantly affected when Drg1 was deleted from cells (Zeng et al., 2021). This observation led us to hypothesize that this GTPase plays a key role in maintaining mitochondrial function. Mitochondria support ATP production, intermediary metabolism, cellular signaling, and stress adaptation. Although mitochondria retain their own genome and translation machinery, mammalian mitochondrial DNA encodes only 13 proteins. Approximately 99% of the mitochondrial proteome is encoded in the nucleus, synthesized by cytoplasmic ribosomes, and subsequently targeted to mitochondria (Rath et al., 2021). Mitochondrial function therefore depends on the coordinated translation, targeting, import, folding, and assembly of proteins produced outside the organelle. Oxidative phosphorylation, mitochondrial translation, protein import, and mitochondrial dynamics require the stoichiometric assembly of multiprotein complexes. Therefore, even modest disturbances in cytosolic protein production can compromise mitochondrial proteostasis and bioenergetic function.

Mitochondria represent a compelling potential target of Drg1-dependent translation. Their proteome is overwhelmingly produced by cytoplasmic ribosomes, a substantial subset of mitochondrial proteins is translated near the outer mitochondrial membrane, and successful protein biogenesis requires coordination among elongation, targeting, import, and folding. These observations raise several unresolved questions. Is Drg1 positioned within the mitochondrial-proximal translation environment? Does Drg1 loss alter the translational engagement or protein output of nuclear-encoded mitochondrial transcripts? If so, are these molecular defects accompanied by changes in mitochondrial architecture and bioenergetic function?

Here, we identify mitochondrial homeostasis as an important physiological output of Drg1-dependent translation. We show that a fraction of the Drg1–Dfrp1 complex associates with the cytosolic face of the mitochondrial outer membrane, where cytoplasmic ribosomes are enriched. Mitochondrial-fraction APEX2 proximity proteomics identifies a Drg1-associated molecular environment containing proteins involved in mitochondrial import, oxidative phosphorylation, mitochondrial translation, and proteostasis. Loss of Drg1 selectively alters ribosome occupancy on nuclear-encoded mitochondrial transcripts and changes protein output from a mitochondria-targeted reporter. These molecular alterations are accompanied by remodeling of the mitochondrial network and reductions in mitochondrial membrane potential and ATP production. Together, our findings support a model in which the ribosome-associated translation factor Drg1 contributes to the production and homeostasis of nuclear-encoded mitochondrial proteins and thereby helps maintain mitochondrial architecture and bioenergetic function.

## Results

### 1. Subcellular localization of Drg1 via immunofluorescence microscopy

We first determined the subcellular localization of Drg1 using immunofluorescence confocal microscopy. In this experiment, the nucleus and mitochondria were stained with the fluorescent dye DAPI and MitoTracker, respectively, and their location in the cell was observed using a confocal microscope. As shown in **Figure 1**, Drg1 protein predominantly localizes in the cytoplasm, consistent with their roles in promoting protein synthesis in the ribosome. Using the same technique, we observed that Dfrp1 is almost exclusively localized in the cytoplasm (**Supplemental Figure S2**).

**Figure 1.**
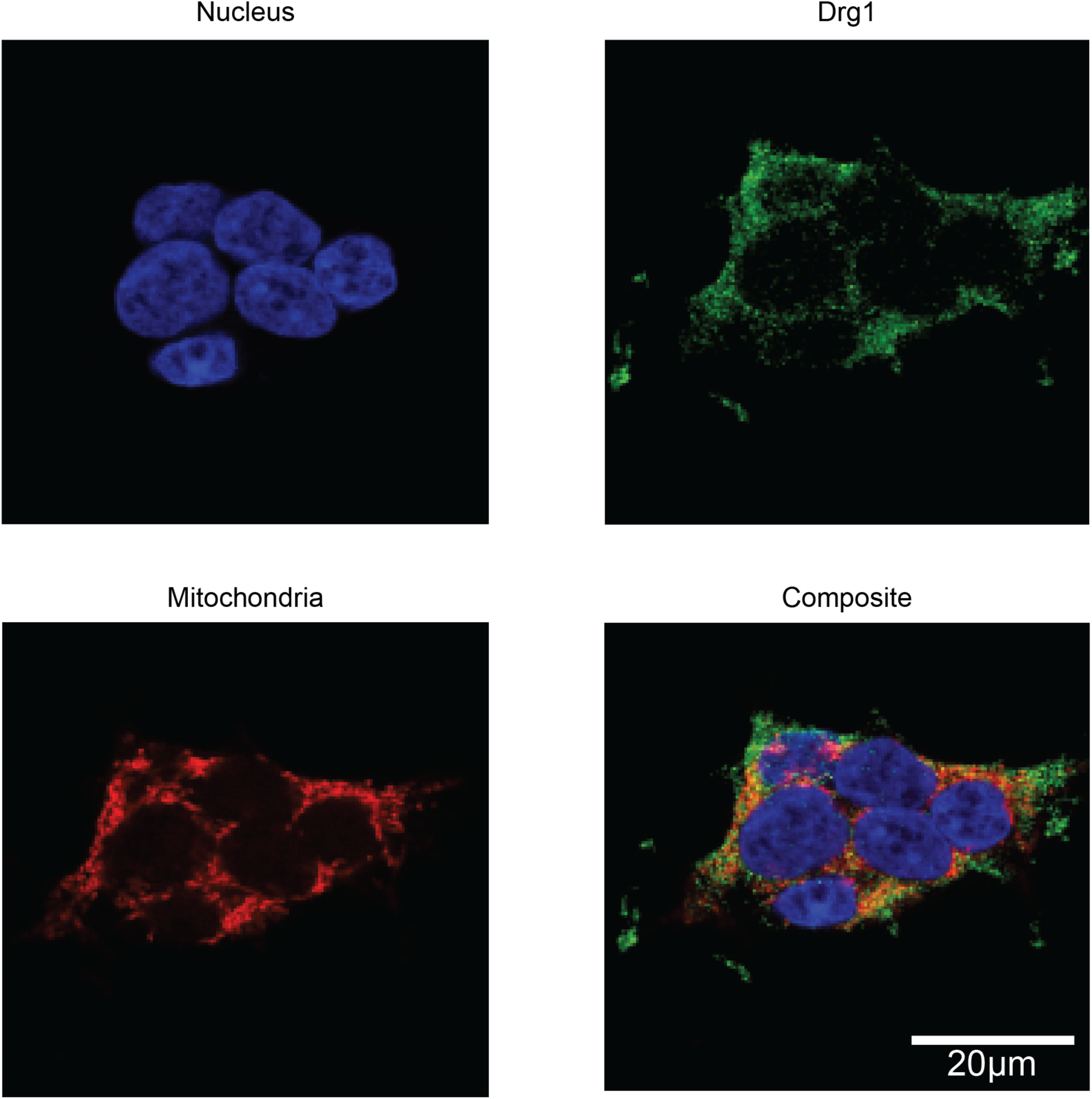
Subcellular localization of Drg1. Immunofluorescence confocal imaging of Drg1. Nuclei were stained with DAPI (blue), mitochondria with MitoTracker (red), and Drg1 with antibodies (green, as indicated). Drg1 proteins predominantly localize to the cytoplasm.

### 2. A fraction of the Drg1/Dfrp1 complex associates with the mitochondrial outer membrane

We next asked whether Drg1 is positioned to influence the synthesis or delivery of nuclear-encoded mitochondrial proteins. We therefore examined the subcellular distribution of endogenous Drg1 using super-resolution Airyscan imaging and biochemical fractionation. First, we examined Drg1 mitochondrial localization using immunofluorescence Airyscan super-resolution imaging. As shown in **Figure 2A**, Drg1 forms a diffuse cytoplasmic pool, with a part of it substantially overlapping with the mitochondrial network. To quantify this overlap, we calculated Pearson’s correlation coefficient between Drg1 and mitochondrial signal across 3 biological replicates and 5 independent fields. This analysis gave us a Pearson’s correlation value ranging between 0.79-0.83 (**Supplemental Figure S3**), indicating that at least a fraction of Drg1 signal spatially coincides with mitochondria, in addition to being present in the cytosol.

**Figure 2.**
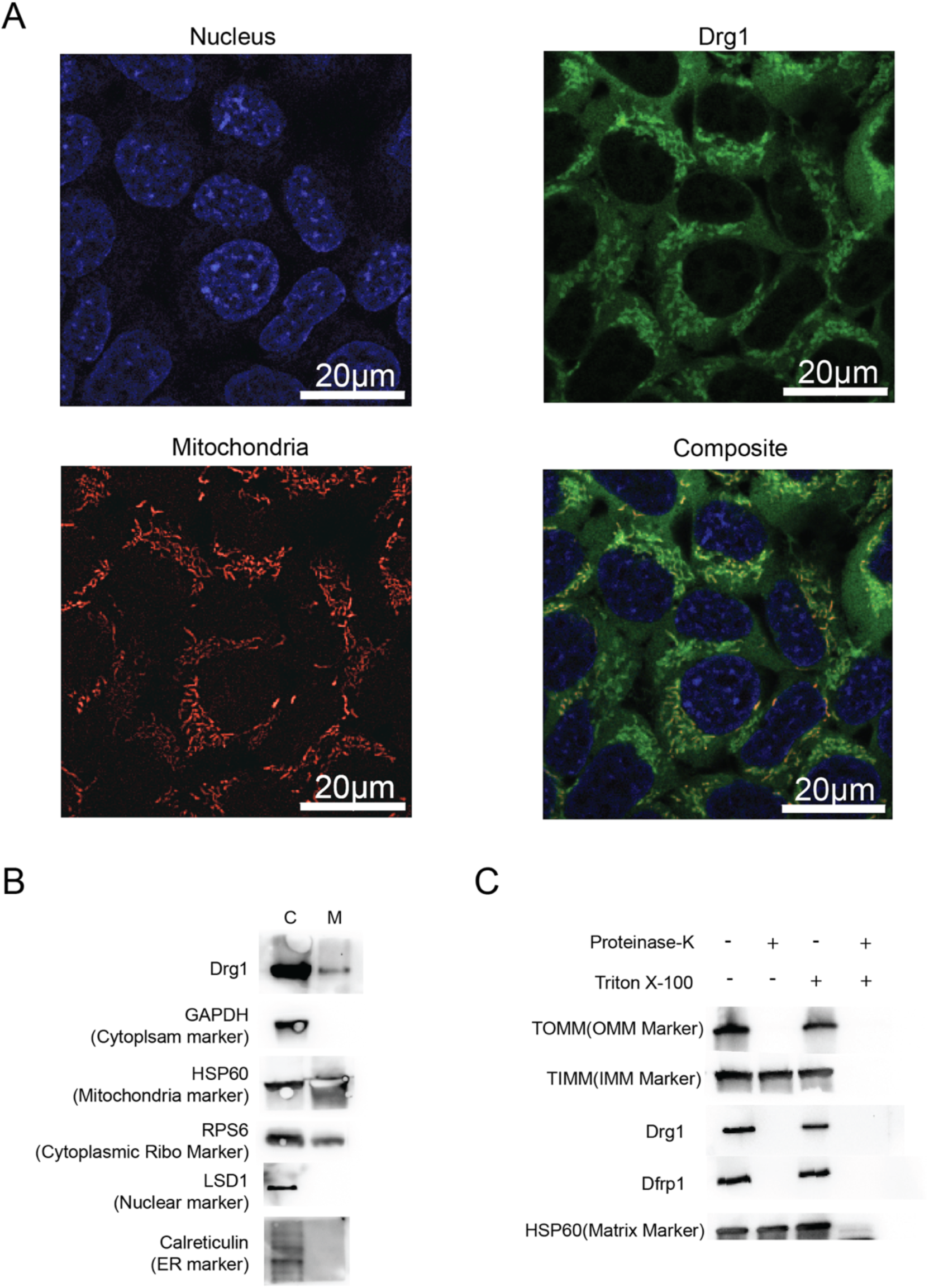
A fraction of Drg1 localizes to the mitochondrial outer membrane. A. Immunofluorescence Airyscan imaging of Drg1 showing mitochondrial localization. Nuclei were stained with DAPI (blue), mitochondria with MitoTracker (red), and Drg1 with antibodies (green, as indicated). Drg1 protein here also looks to predominantly localize to the cytoplasm however a portion of it, is shown to localize to the mitochondria. The Pearson’s correlation calculation is shown in **Supplemental** Figure 3. B. Biochemical fractionation shows Drg1 presence in cytoplasmic and mitochondrial fractions. HEK293T cells were lysed and fractionated to mitochondrial and cytoplasmic fractions and were probed for the presence of Drg1 and RPS6 in the mitochondrial fraction using Western Blot. GAPDH, HSP60, LSD1, and Calreticulin serve as control markers for purity of extraction. Data show Drg1 and RPS6(cytoplasmic ribosome marker) localize to mitochondria. C. Controlled protease protection assay using Proteinase K on purified mitochondria, demonstrating that Drg1 and Dfrp1 are localized to the outer mitochondrial membrane (OMM).

Second, we confirmed Drg1 mitochondrial localization using biochemical methods. In this experiment, we isolated mitochondria using the differential centrifugation method as described (Senichkin et al., 2021). The presence of Drg1 was confirmed in both cytoplasmic (C) and mitochondrial (M) fraction by their respective antibodies (see Materials and Methods section). Fraction purity was confirmed by using GAPDH (cytoplasmic marker), LSD1(Nuclear marker), and Calreticulin (ER marker) being absent from the mitochondrial fraction, with only HSP60 (mitochondrial marker) being present in the mitochondrial fraction, as shown in **Figure 2B**. Furthermore, Dfrp1 was also confirmed to be found in the cytoplasmic and mitochondrial fractions by their respective antibodies (see Materials and Methods section) using Western Blot. As shown in **Supplemental Figure S4**, in wild-type (WT) cells, both Drg1 and Dfrp1 were present in the cytoplasmic and mitochondrial fractions. By contrast, using the ΔDrg1 HEK293T cell as a comparison, no Drg1 was observed in the mitochondrial fraction, as expected. Of note, little Dfrp1 was shown in the mitochondrial fraction. These observations indicate that Dfrp1’s mitochondrial localization is significantly affected in the absence of Drg1. Thus, it is highly likely that the two proteins, Drg1 and Dfrp1, bind as a heterodimeric protein complex to associate with the mitochondria in the cytoplasm.

Mitochondria are separated from the cytoplasm by the outer and inner mitochondrial membranes (OMM and IMM). Since neither Drg nor Dfrp contains mitochondrial localization signal sequences in their amino acid sequences (**Supplemental Figure S1)**, we predict that the protein complex only associates with the outer mitochondrial membrane, instead of residing inside mitochondria. To test this prediction, we isolated mitochondria using controlled detergent-proteinase treatment and separated the mitochondrial inner and outer mitochondrial proteins. In this experiment, the native mitochondria were purified by differential centrifugation without adding any detergent to hypotonic and hypertonic lysis buffer (Senichkin et al., 2021) and subsequently underwent Triton X-100 and proteinase K treatment (Zhou et al., 2023). The presence of 10 µg of proteinase K per mg of mitochondrial protein allows for partial digestion of proteins bound to the OMM only and leaves proteins bound to the IMM and mito-matrix intact. However, the addition of both Triton X-100 and proteinase K simultaneously dissolves the mito-lipid bilayer and digests all proteins in the mitochondria. Using the known mitochondrial proteins TOM40, TIM29 and HSP60 as a marker for OMM-bound, IMM-bound, and mitochondrial matrix proteins, respectively, we thus probed the sub-mitochondrial localization of the Drg complex. As shown in **Figure 2C**, addition of proteinase K in the absence of Triton X-100 destroyed the Drg/Dfrp complex, as well as the OMM-bound protein marker TOM40, leaving only proteins bound to IMM and mito-matrix to be detected. By contrast, the same complex is present in the purified mitochondria when proteinase K treatment is omitted. This result strongly supports the binding of the Drg complex to the outer membrane of mitochondria.

Together, the imaging, fractionation, and protease-accessibility results demonstrate that Drg1 is predominantly cytoplasmic but that a detectable fraction of the Drg1/Dfrp1 complex associates with the cytosolic face of the mitochondrial outer membrane. We next asked whether this localization coincides with the presence of cytoplasmic ribosomes at the mitochondrial surface.

### 3. Cytoplasmic ribosomes are enriched near the mitochondrial surface

Many nuclear-encoded mitochondrial proteins are synthesized by cytoplasmic ribosomes positioned near mitochondria (Luo et al., 2025; Weis et al., 2013; Zhu et al., 2025). To determine whether Drg1 occupies such a spatial environment, we examined the distribution of Drg1-bound cytoplasmic ribosomes relative to the mitochondrial network.

In this investigation, immunofluorescence Airyscan super-resolution microscopy was performed using the small ribosomal subunit protein Rps6, and MitoTracker to label ribosomes and mitochondria, respectively. These images revealed the proximity of ribosomes around mitochondria (**Figure 3A**). To quantify this relationship independent of a fixed colocalization threshold, we measured RPS6 intensity as a function of distance from the mitochondrial surface, binning pixels into concentric 100 nm shells extending outward from a mitochondrial mask (**Supplemental Figure S5A** and **B**). RPS6 signal was highest immediately adjacent to mitochondria and declined progressively with increasing distance, from a mean intensity of ∼3.0 fold within 100 nm of the mitochondrial surface to background cytoplasmic levels (∼1.0) by 1900–2000 nm (**Supplemental Figure S5C**). This distance-dependent enrichment supports a genuine spatial association between cytoplasmic ribosomes and mitochondria, rather than an artifact of overlapping signal in densely packed cytoplasm. Consistent with the imaging results, western blot of cellular fractions demonstrated that Rps6 and Drg1 are present in the mitochondrial fraction (**Figure 2B** and **3B**), supporting the imaging results. The enrichment of ribosomal maker signal near mitochondria, together with the recovery of ribosomal proteins in mitochondrial fractions, supports the presence of a mitochondrial-proximal cytoplasmic translation environment. We therefore examined whether the ribosome-interacting region of Drg1 contributes to its mitochondrial association.

**Figure 3.**
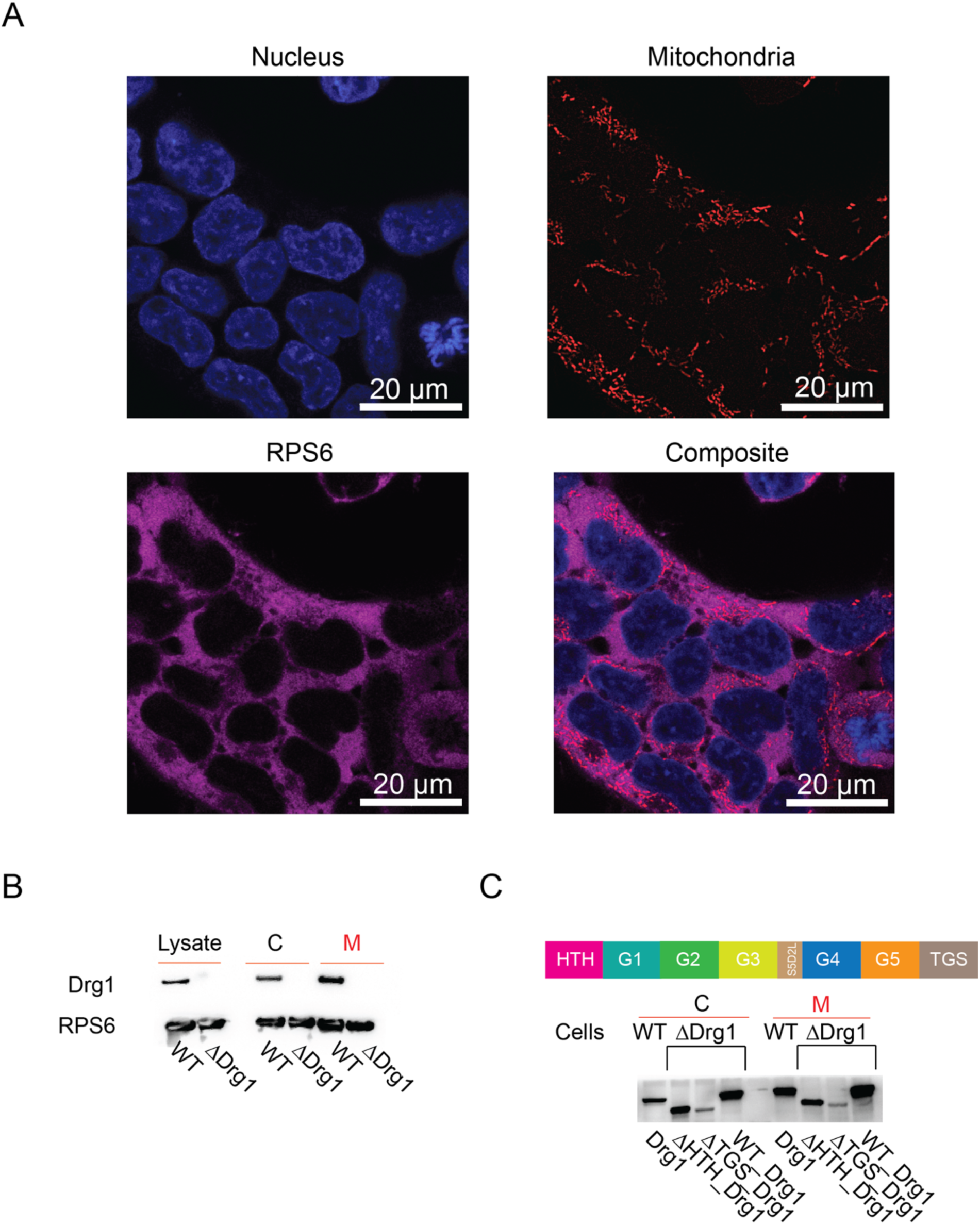
Drg1 associates with cytoplasmic ribosomes at the mitochondrial outer membrane. A. Immunofluorescence microscopy showing cytoplasmic ribosomes (shown via its marker Rps6, purple) with partial colocalization at mitochondria (MitoTracker, red); nuclei are stained with DAPI (blue). Cytoplasmic ribosomes exhibit enrichment at the mitochondrial periphery. See Supplemental Figure 5 for calculations. B. Western blot analysis of cytoplasmic (C) and mitochondrial (M) fractions confirms the presence of both Drg1 and cytoplasmic ribosomes in the mitochondrial fraction. Ribosomes were detected via Rps6 marker. Note that deletion of cellular Drg1 has little effect on the presence of ribosomes. Drg1 localization at the mitochondrial OMM depends on its own expression. C. The TGS domain of Drg1 is required for its association with cytoplasmic ribosomes at the mitochondrial outer membrane. The WB was performed by loading equal amount of proteins quantified by Bradford assay. Full-length Drg1 and ΔHTH_Drg1 localize to the mitochondrial fraction in ΔDrg1 HEK293T cells, whereas ΔTGS_Drg1 fails to do so. Because the TGS domain is specifically required for ribosome and Dfrp1 binding, but not microtubule association, these results indicate that Drg1 localizes to mitochondria through interaction with ribosome-associated complexes rather than microtubules.

### 4. The ribosome-binding TGS domain contributes to Drg1 association with mitochondria

To test whether ribosome interaction contributes to the mitochondrial association of Drg1, we compared the mitochondrial recovery of wild-type Drg1 with that of domain-deletion variants. Since Drg1 was previously reported to bind microtubules and promote their dynamics *in vitro* (Schellhaus et al., 2017), and microtubules play a central role in mitochondrial dynamics—including fusion, fission, and motility (Detmer et al., 2007; Westermann and 2008) as mitochondria walk towards each other along microtubules during the fusion process, and divide and move away from each other along microtubules during fission, we sought to exclude the possibility that Drg1’s apparent mitochondrial OMM association occurs indirectly via microtubules. Prior work showed that both the HTH and TGS domains of Drg1 are required for microtubule binding and promotion of tubulin polymerization (Schellhaus et al., 2017), whereas only the TGS domain is required for Drg1/Dfrp1 complex formation (Francis et al., 2012), which indispensably mediates ribosome association.

To distinguish between these possibilities, we deleted either the HTH or TGS domain of Drg1, and generated ΔHTH_Drg1 and ΔTGS_Drg1 mutants. We then transfected these mutant proteins into ΔDrg1 cells and examined their mitochondrial localization in the cell. Mitochondrial fractions from ΔDrg1 cells transfected with these mutant constructs were analyzed by quantitative Western blot. As shown in **Figure 3C** and **Supplemental Figure S6**, deletion of the TGS domain—but not the HTH domain—diminished Drg1 association with mitochondria. Wild-type Drg1 (WT_Drg1) was also used as a control in this experiment simultaneously.

Of note, deletion of the TGS domain reduced Drg1 recovery in the mitochondrial fraction, whereas deletion of the HTH domain had a smaller effect. The pattern is similar to the protein’s association with translating ribosome on domain deletion (Francis et al., 2012). These findings indicate that the TGS domain contributes to mitochondrial association, consistent with—but not by itself proving—a model in which a fraction of Drg1 reaches the mitochondrial surface in association with cytoplasmic ribosomes. We next characterized the protein environment surrounding mitochondrial-associated Drg1.

### 5. Mitochondrial-fraction Drg1–APEX2 proteomics identifies proteins involved in mitochondrial translation, import, and oxidative phosphorylation

Having independently detected Drg1 at the mitochondrial outer membrane, we used APEX2 proximity labeling (Lam et al., 2015) to characterize proteins recovered within its mitochondrial-proximal environment. In this experiment, wild-type HEK293T cells were transfected with a plasmid expressing Drg1 fused to the engineered soybean ascorbate peroxidase APEX2 and a V5 epitope tag (**Supplemental Figure S7 A-C**). Upon addition of biotin–phenol and hydrogen peroxide, APEX2 catalyzed the biotinylation of proteins located within approximately 20 nm of Drg1. Biotinylated proteins were captured by streptavidin beads and analyzed from whole-cell lysates, polysome-associated fractions isolated by sucrose-gradient fractionation, and purified mitochondrial fractions by mass spectrometry–based proteomic analysis with three biological replicates analyzed for each condition.

Assuming Drg1’s mitochondria association is via translating ribosomes, we compared polysome versus mitochondrial fractions in APEX2-based proximity proteomics analysis, we observed a predominance of proteins annotated as nuclear, with mitochondrial proteins comprising a comparable fraction and a smaller subset annotated as ER (**Supplemental Figure S8 and Supplemental Table 1**). This distribution likely reflects the spatial organization of intracellular translation, as mitochondria frequently localize in close proximity to both the nucleus and ER, creating shared translational microenvironments. In addition, the ∼20 nm effective labeling radius of APEX2 proximity tagging can capture proteins translated near the mitochondrial surface even if their canonical steady-state localization is close to the nucleus or ER.

Within the mitochondrial fraction, the enriched mitochondrial proteins were strongly biased toward matrix and inner mitochondrial membrane localization (**Table 1**). Among the significantly enriched hits are mitochondrial respiratory chain components and mito-ribosomal proteins (for example, ATPIF1, PYCR2, NDUFS4, MRPS5, MRPS30). The data also show enrichment of TOMM70A, a receptor on the outer membrane of mitochondria that recognizes precursor proteins with mitochondrial targeting sequences. Cytoplasmic ribosomal proteins, such as RPL32, RPL13, RPL10, RPS15, and RPS17, were also identified in the dataset. Since we compared mito-fraction versus polysome-fraction, most cytoplasmic ribosomal proteins were neither significantly enriched nor depleted, as expected due to our spatial comparison of mitochondrial versus ribosomal fractions in the APEX2 design. Thus, these results suggest that Drg1 localizes to regions of translation at the mitochondrial surface, consistent with a model in which nascent chains bearing N-terminal mitochondrial targeting sequences are translated at or near the mitochondrial outer membrane (OMM), engaging in localized translation and subsequent import.

**Table 1.**
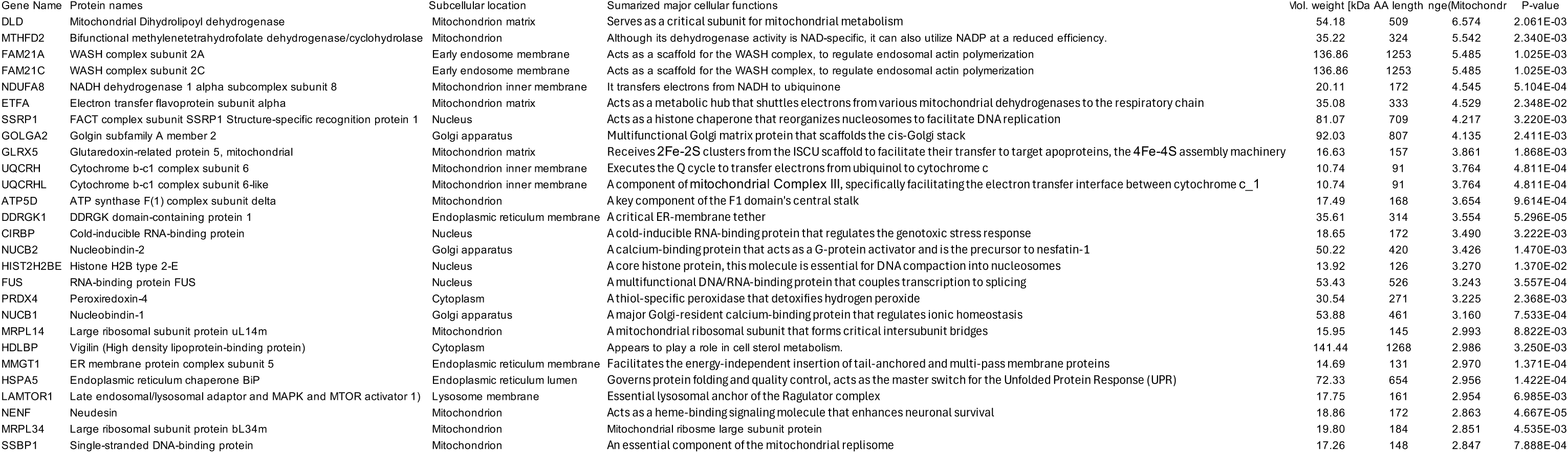
Top 27 proteins enriched in the mitochondrial fraction of the Drg1 interactome identified by APEX2-based proximity labeling proteomics. Data represent the top 27 proteins ranked by fold change from a total of 154 significantly enriched proteins identified in the Drg1 interactome, comparing mitochondrial and polysome fractions (log2FoldChange [Mitochondria/Polysome] > 1; *Pvalue* < 0.05). In addition to mitochondrial proteins, hits include proteins of nuclear and endoplasmic reticulum origin, reflecting the broader subcellular distribution of Drg1-interacting partners. Columns indicate gene name, protein name, subcellular location, function, molecular weight (kDa), sequence length (amino acids), log₂2FoldChange (Mitochondria/Polysome), and *P*-value.

Notably, the Drg1-associated mitochondrial proteins are enriched for electron transport chain components—particularly ATP synthase subunits—as well as proton-coupled transporters, a subset of mitochondrial ribosomal proteins, and a limited number of mitochondrial metabolic enzymes. These functional classes are central to mitochondrial ATP production, membrane potential maintenance, and proteostasis, aligning with the mitochondrial morphological and bioenergetic defects we observe upon Drg1 loss discussed later (**Section 9**). Importantly, many of the identified mitochondrial proteins overlap with nuclear-encoded mitochondrial proteins that undergo localized translation prior to mitochondrial import (Rath et al., 2021). Protein names shown in bold in our datasets have been independently validated as locally translated at OMM and imported into mitochondria (Luo et al., 2025; Zhu et al., 2025), suggesting that Drg1 may promote localized translation of a subset of nuclear-encoded mitochondrial mRNAs at the mitochondrial outer membrane. Notably, such a function does not seem to be restricted to the coding sequence length, as Drg1 facilitates translation of both long (>∼350–400 amino acids) and short (<∼200 amino acids) nuclear-encoded mitochondrial proteins (**Table 1**). Together, these results support a model in which Drg1 promotes efficient localized translation at the OMM, thereby protecting mitochondrial function and homeostasis.

The mitochondrial-fraction Drg1–APEX2 dataset contained proteins involved in mitochondrial protein import, oxidative phosphorylation, and mitochondrial translation, including TOMM70A, respiratory-chain components, and mitoribosomal proteins. Although this analysis does not establish direct binding or Drg1-dependent synthesis of individual proteins, it identifies mitochondrial pathways and candidate transcripts potentially influenced by Drg1-associated translation. We therefore tested whether loss of Drg1 affects mitochondria-targeted translation, as well as the translational engagement of selected mitochondrial protein–encoding mRNAs.

### 6. Loss of Drg1 alters protein output from a mitochondria-targeted reporter

Nuclear-encoded mitochondrial proteins that are cytosolically translated contain N-terminal Mitochondrial Localization/Targeting Signals (MLS/MTS), which are essential for targeting the protein to mitochondria. After entering the mitochondria, the nascent peptide is processed into functional proteins. To test directly whether Drg1 affects translation of a transcript known to localize at the mitochondrial surface, we built a reporter by adding an MLS nucleotide sequence of the human SOD2 gene to the 5’ end of GFP and the 3’UTR of SOD2 (which facilitates mito-surface mRNA localization) after the stop codon of GFP (**Figure 4A, i**), then cloned this construct into a pcDNA plasmid backbone under a CMV promoter (Eliyahu et al., 2023; Kaltimbacher et al., 2006; Michaelis et al., 2022). Reporter output was compared between wild-type and ΔDrg1 cells using single-and dual-fluorescence designs.

**Figure 4.**
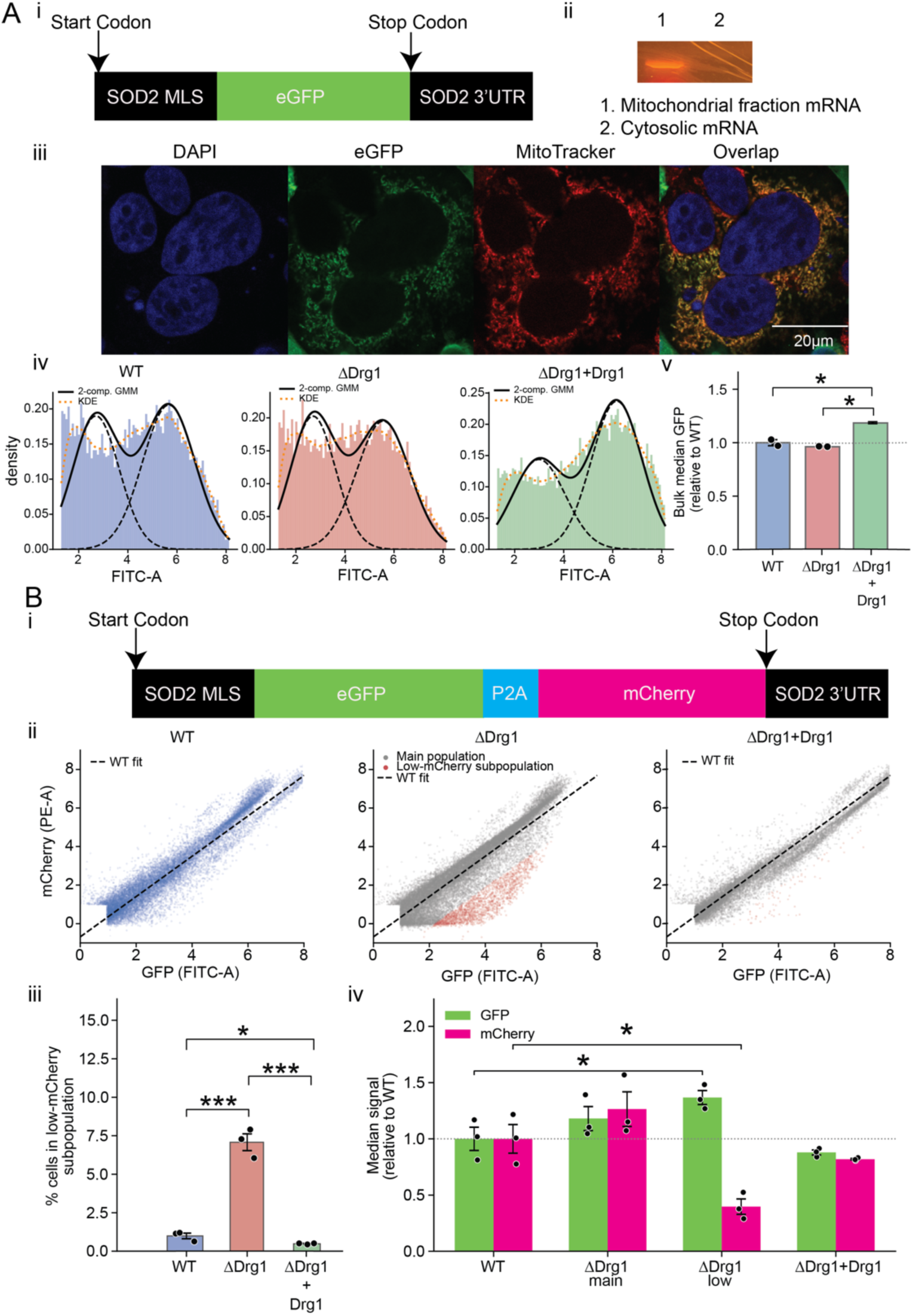
**Loss of Drg1 impairs localized translation on mitochondria. A. Investigating localized translation on mitochondria using a single eGFP reporter.** Schematic of a mitochondrial import reporter consisting of the SOD2 mitochondrial targeting sequence (MTS) and SOD2 3′UTR flanking an eGFP coding region. SOD2 is a nuclear-encoded mitochondrial enzyme (mitochondrial peroxidase), and the MTS/3′UTR elements promote delivery of newly synthesized eGFP to mitochondria after transfection into HEK293T cells. Import efficiency is quantified by eGFP fluorescence. (ii) Localization of the eGFP mRNA to the mitochondria. The DNA construct shown in (A) was cloned and transfected into HEK293T cells. Following mitochondrial and cytosolic fractionation, translating ribosomes were pelleted and mRNA was extracted. RT-PCR analysis confirms the presence of construct mRNA in the mitochondrial polysome fraction. (iii). Functional eGFP proteins are only found in mitochondria. HEK293T cells were transfected with the plasmid containing the cloned DNA construct. Fluorescence confocal imaging reveals eGFP (green) localizing to mitochondria that were labeled with MitoTracker (red). Nuclei were counterstained with DAPI (blue). (iv) and (v) A reduced median of eGFP fluorescence signal in ΔDrg1 cells. A reduced median of eGFP fluorescence signal in ΔDrg1 cells expressing the reporter compared to WT cells, and the eGFP signal was recovered upon expressing back WT Drg1 into ΔDrg1 cells, as measured by flow cytometry. This observation is consistent with decreased translation capacity in the absence of Drg1. **B. Investigating localized translation on mitochondria using a dual eGFP and mCherry reporter.** (i) Schematic of a mitochondrial import reporter consisting of the SOD2 mitochondrial targeting sequence (MTS) and SOD2 3′UTR flanking an eGFP-P2A-mCherry coding region. SOD2 is a nuclear-encoded mitochondrial enzyme (mitochondrial peroxidase), and the 3′UTR elements promote delivery of newly synthesized mRNA to the mitochondrial OMM. During translation, presence of the MTS shuttles eGFP to mitochondria, while mCherry will be cytosolic after P2A ribosomal skipping. This construct was cloned into a pcDNA vector and was transfected into HEK293T cells. (ii) A reduced median of mCherry fluorescence signal in ΔDrg1 cells. Representative Flow cytometry scatter plots of mCherry versus GFP signal in WT, ΔDrg1 cells, as well as ΔDrg1 cells with WT Drg1 supplemented; a discrete low-mCherry subpopulation (red) is evident in ΔDrg1 cells relative to the WT fit (dashed line). The low-mCherry intensity was recovered upon supplementing WT Drg1 protein into the ΔDrg1 cells. (iii) Quantification of low mCherry cell population and (iv) Quantification of median GFP and mCherry signal in the dual reporter assay. Median GFP and mCherry intensities relative to WT was quantified after fitting gaussian bimodal distribution to the raw data in the cell populations of WT, ΔDrg1 main (grey in ii), and ΔDrg1 low-mCherry (red in ii), ΔDrg1 cells with WT Drg1 supplemented.

We first tested and confirmed that mRNA and the protein localize to mitochondria (**Figure 4A, ii and iii**). Then, we transfected WT and ΔDrg1 HEK293T cells with this construct and used flow cytometry to determine the median GFP intensity, indicative of functional GFP protein expression in mitochondria in the cell (**Figure 4A-iv**). As shown in **Figure 4A-v**, the median GFP intensity in ΔDrg1 cells is reduced ∼5% compared to WT cells, and such a reduction was rescued upon expressing WT Drg1 into the ΔDrg1 cells.

The observed decrease in GFP intensity is likely due to impaired mitochondrial protein import, reduced translation or reporter mRNA stability. To distinguish impaired mitochondrial import from altered translation of the SOD2 reporter, we generated a dual-reporter construct in which SOD2-targeted eGFP was followed by a P2A ribosomal-skipping sequence and cytosolic mCherry, with expression controlled by the same SOD2 3′UTR (**Figure 4B-i**). Translation through the P2A sequence produces two physically separate reporter proteins from the same mRNA and the same ribosomal translation event. Thus, a uniform reduction in transcript abundance or translation initiation would be expected to decrease both eGFP and mCherry. Correct targeting and folding of the GFP was confirmed in a similar way as described earlier in **Figure 4A-ii**. In the major cell population, both eGFP and mCherry fluorescence in ΔDrg1 cells were comparable to WT (**Figure 4B-ii**). Strikingly, however, a distinct population characterized by low mCherry fluorescence was observed almost exclusively in ΔDrg1 cells; this subpopulation was identified by fitting a two-component Gaussian model to the mCherry residuals relative to the WT fit. Within this population, mCherry fluorescence decreased to ∼30% of WT (P = 0.010), whereas eGFP fluorescence was modestly increased. Addition of the WT Drg1 back into the ΔDrg1 cells restores the eGFP/mCherry ratio, producing a profile similar to that observed in the WT cells (Figure 4B-ii, iii, iv). The discordant behavior of the two reporters is difficult to reconcile with a simple reduction in transcription, mRNA abundance, or translation initiation of the reporter, which would be expected to affect both products in the same direction.

Instead, the uncoupling of upstream eGFP and downstream mCherry suggests that loss of Drg1 affects co-translational mRNA translation and stability. Of note, changes in mRNA stability likely reflects changes in mRNA translation, as better-translated mRNA is more stable. Thus, Drg1 may preferentially affect the translation of the 3’-end of the coding sequence. Additionally, altered translation elongation or processivity, or even differential maturation or stability of the reporter proteins, could also be an alternative explanation. More future investigations are required to distinguish among these possibilities. Nevertheless, our data show that loss of Drg1 perturbs protein synthesis associated with the mitochondrial outer membrane. Importantly, the dual-reporter phenotype is restricted to a subpopulation of ΔDrg1 cells, suggesting that the translational consequences of Drg1 loss are heterogeneous and may depend on cell-to-cell differences in translational state or mRNA fate.

### 7. Loss of Drg1 selectively alters ribosome occupancy on mitochondrial protein–encoding mRNAs

As a cytoplasmic GTPase, Drg1 does not localize inside mitochondria and has no known mitochondria-specific biochemical functions. We therefore infer that the mitochondrial defects observed upon Drg1 loss arise indirectly rather than from a direct role within the organelle. Such a translation defect will conceivably disrupt global translational homeostasis, an underlying molecular basis for the compromised mitochondrial function observed in this study. Mitochondria are highly dynamic organelles that perform numerous essential cellular functions beyond energy production (Shen et al., 2022). Perturbations in mitochondrial function can, in turn, influence cellular protein synthesis and broader physiological processes, thereby establishing a potential feedback loop that further amplifies disruptions in homeostasis. To better understand these interconnected effects, we next employed candidate approaches to assess alterations in cellular homeostasis in the absence of Drg1.

To determine whether Drg1 affects endogenous mitochondrial protein–encoding transcripts, we compared their abundance in total and ribosome-associated mRNA fractions from wild-type and ΔDrg1 cells. Candidate transcripts were selected from mitochondrial pathways represented in the APEX2 dataset and included factors involved in mitochondrial translation, respiration, and proteostasis. In this experiment, we purified total cellular mRNA and ribosome-protected (translating) mRNA from WT and ΔDrg1 HEK293T cells, converted each fraction into cDNA libraries, and quantified selected transcripts by qPCR. We examined mRNAs encoding mitochondrial ribosomal proteins involved in mitochondrial translation (MRPS30 and MRPL43) and ATP5D, a subunit of electron transport chain complex V required for ATP production, as well as other proteins labeled in **Table 1** that are known to be locally translated at mitochondrial OMM. GAPDH served as a normalization control. Threshold cycle (Ct) values for GAPDH were comparable between WT and ΔDrg1 cells in both total and translating mRNA pools, confirming equivalent input and fractionation efficiency. No significant differences were observed in the total mRNA levels of genes in **Figure 5**. In contrast, ribosome occupancy—defined as the ratio of transcript abundance in the translating mRNA pool to that in the total mRNA pool—was substantially altered for each gene in ΔDrg1 cells, ribosomal occupancy decreased for seven transcripts (MRPS30, ATP5D, ATP5I, NDUFB1, NDUFAF2, ATP5F1, and ETFA) and increased for four (MRPL43, NDUFS4, CLPP, and UQCRQ), indicating altered translational engagement upon loss of Drg1 in a transcript-specific way (**Figure 5**).

**Figure 5.**
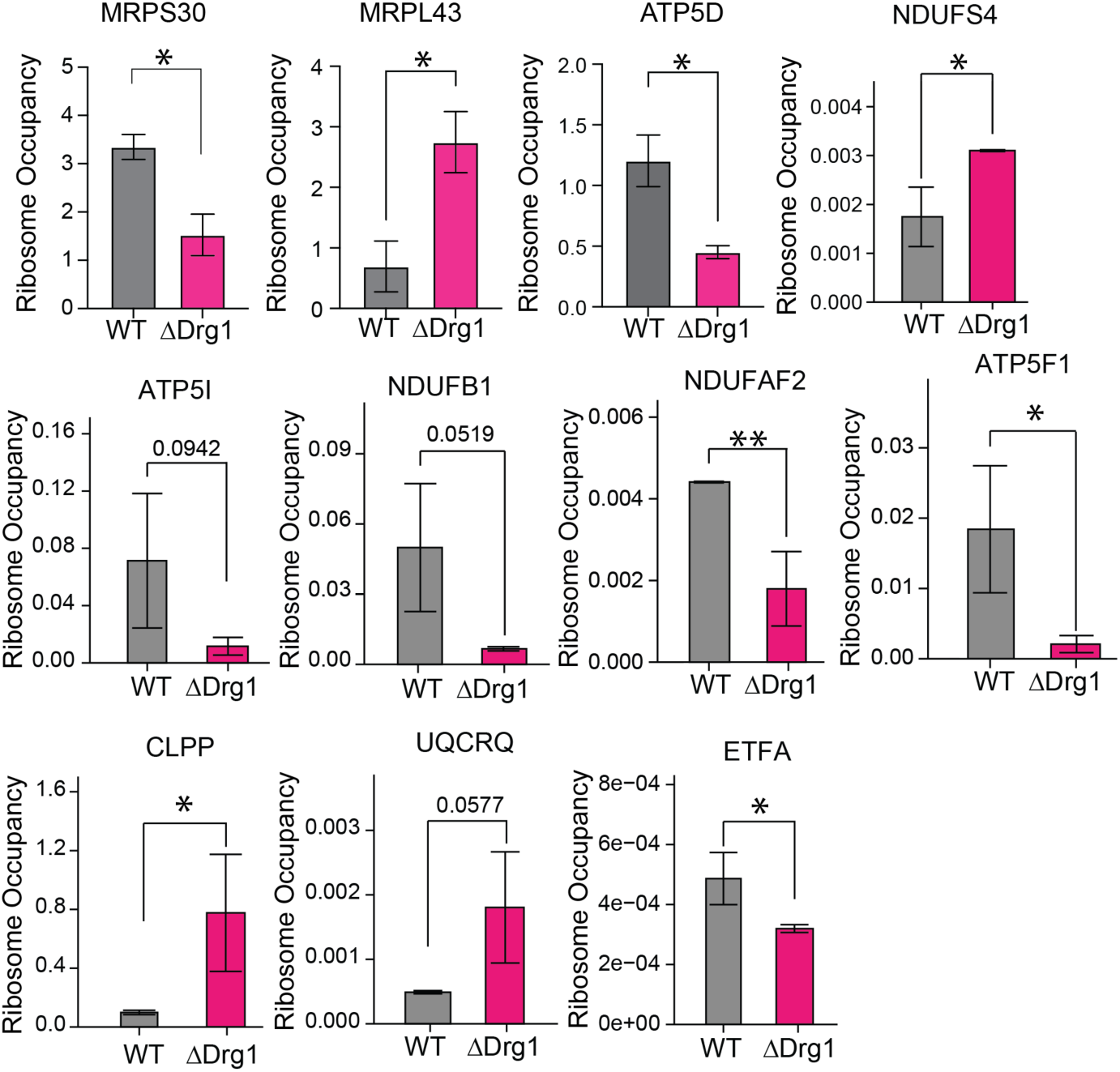
Loss of Drg1 compromises cellular protein homeostasis. Loss of Drg1 alters degree of ribosome occupancy of mitochondrial protein–encoding mRNAs without changing total mRNA abundance. Total cellular mRNA and ribosome-protected (translating) mRNA were purified from WT and ΔDrg1 HEK293T cells, converted into cDNA libraries, and analyzed by qPCR. Transcripts encoding mitochondrial ribosomal proteins involved in mitochondrial translation (MRPS30 and MRPL43) and ATP5D, a subunit of the electron transport chain complex V required for ATP production, and several other hits mentioned in Figure 5 (NDUFS4, NDUFB1, NDUFAF2, ATP5F1, CLPP, ETFA, UQCRQ, etc.) were examined. GAPDH served as a normalization control. Ct values for GAPDH were comparable between WT and ΔDrg1 cells in both total and translating mRNA pools, indicating equivalent input and fractionation efficiency. No significant differences were detected in total mRNA levels between WT and ΔDrg1 cells (data not shown). In contrast, ribosome occupancy—calculated as the ratio of transcript abundance in the translating fraction to that in the total mRNA fraction—was substantially altered for each gene in ΔDrg1 cells, indicating impaired translational engagement of mitochondria-related mRNAs upon loss of Drg1.

Total abundance remained largely unchanged for the examined mRNAs, whereas their ribosome occupancy was altered in a transcript-dependent manner following Drg1 loss. Thus, Drg1 is not required uniformly for mitochondrial mRNA expression but contributes to the translational state of a defined subset of nuclear-encoded mitochondrial transcripts. The endogenous ribosome-occupancy measurements, together with the results obtained from mitochondria-targeted reporter in the previous section, indicate that loss of Drg1 perturbs translation of proteins functionally important to mitochondria. We next determined whether these molecular changes are accompanied by defects in mitochondrial organization.

### 8. Drg1 loss disrupts mitochondrial network organization

Mitochondria distribution within cells is dynamic, and their activity impacts nearly all aspects of cell physiology. Mitochondrial architecture depends on the coordinated synthesis, import, and assembly of nuclear-encoded proteins. Therefore, we examined whether loss of Drg1 alters mitochondrial morphology and network organization. In this experiment, we examined mitochondrial morphology in WT and ΔDrg1 cells using Airyscan Fluorescence microscopy (**Figure 6A**). Our data revealed that ΔDrg1 cells contained fewer, larger mitochondria than WT: mean mitochondrial area increased from ∼1.2 µm² to ∼1.7 µm² (**Figure 6B**), while mitochondrial count per field decreased by ∼20% (**Figure 6B**). Form factor and branch junctions per mitochondrion were both modestly but significantly increased in ΔDrg1 cells (**Figure 6B**), indicating a more interconnected network rather than simple swelling. Aspect ratio was unchanged (WT vs. ΔDrg1, p = 0.68; **Figure 6B**), indicating individual mitochondria were not more elongated or rounded in ΔDrg1 cells. This phenotype suggests network remodeling of mitochondria, not a change in organelle shape.

**Figure 6.**
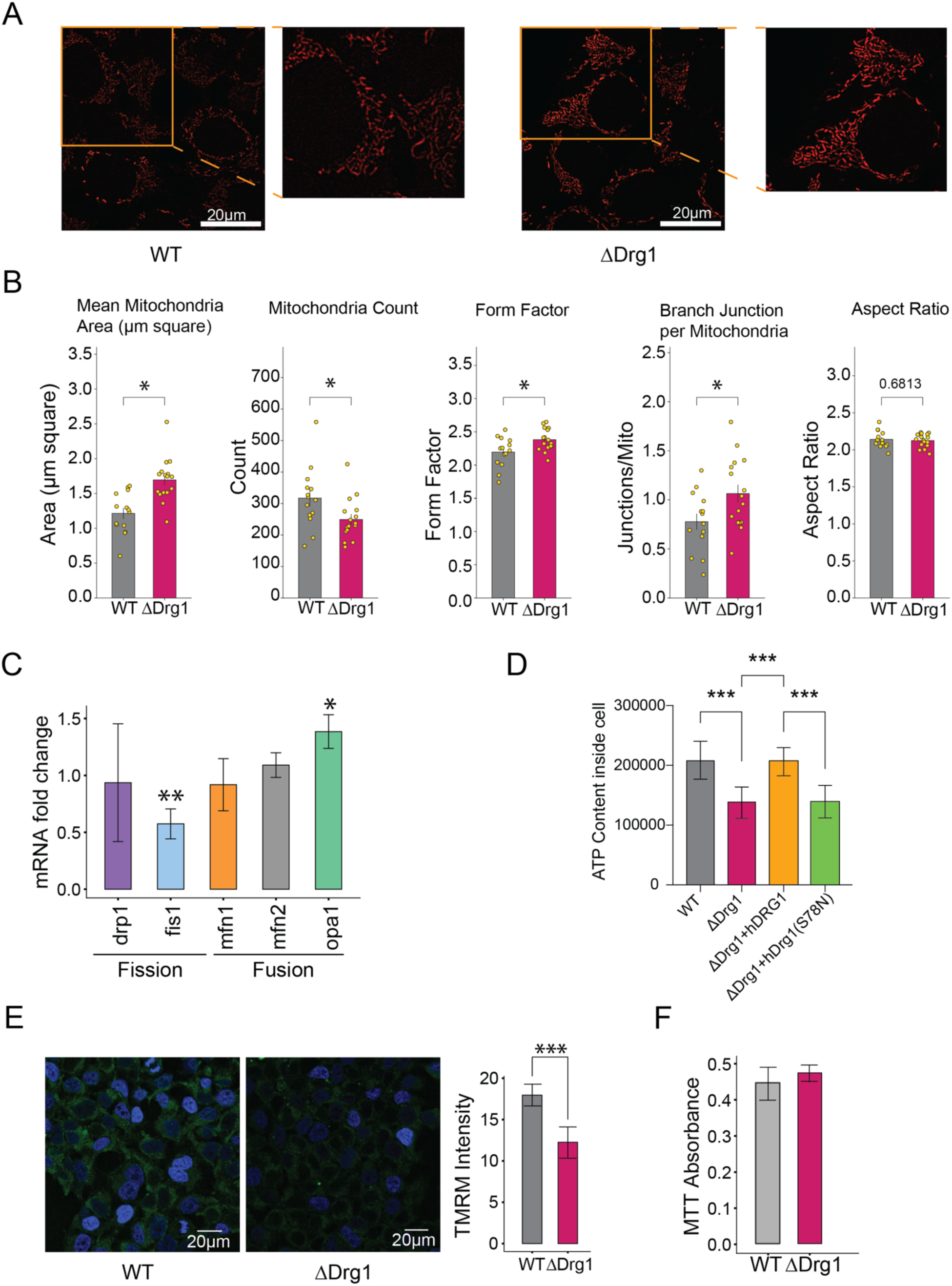
Loss of Drg1 alters mitochondrial morphology and impairs mitochondrial function. (A-C) Loss of Drg1 affects mitochondrial network and architecture. A Representative Airyscan super-resolution images of the mitochondrial network in WT and ΔDrg1 HEK293T cells, with magnified insets. B. Quantification of mitochondrial morphology in WT versus ΔDrg1 cells (individual fields and biological replicates shown as points, mean ± SEM): mean mitochondrial area, mitochondrial count, form factor, branch junctions per mitochondrion, and aspect ratio. ΔDrg1 mitochondria are larger, fewer in number [in the field of view], and more branched, with increased form factor and junctions per mitochondrion; aspect ratio is unchanged C. Loss of Drg1 alters the mRNA abundance of genes involved in mitochondrial fusion and fission, as quantified by qPCR. D. Relative cellular ATP levels in WT, ΔDrg1 cells, as well as ΔDrg1 cells with WT Drg1 or Drg1(S78N mutant) expressed. Loss of Drg1 reduces ATP levels, and the phenotype was rescued upon expressing WT Drg1 back into the ΔDrg1 cells, whereas the GTPase-deficient mutant, Drg1(S78N), failed to rescue. In this experiment, 100ng of plasmid DNA containing WT and mutant human Drg1 (S78N) was transfected into ∼10,000 cells. E. Reduced mitochondrial membrane potential in ΔDrg1 cells. Reduced mitochondrial membrane potential in ΔDrg1 cells compared to WT, measured by TMRM staining (in green) and fluorescence confocal microscopy imaging. F. Loss of Drg1 alone does not induce apoptosis. MTT assay measuring cellular viability and proliferation shows no significant difference between WT and ΔDrg1 cells despite changes in mitochondrial morphology and function.

The altered mitochondrial morphology observed in ΔDrg1 cells indicates disruption of mitochondrial dynamics, which depend on the balanced regulation of fusion and fission as well as proper organelle distribution. Mitochondrial fusion involves coordinated outer and inner membrane fusion mediated primarily by the OMM proteins mitofusin 1 and 2 (Mfn1 and Mfn2) and the IMM protein optic atrophy 1 (OPA1) (Cipolat et al., 2004; Ishihara et al., 2004), whereas mitochondrial fission is controlled largely by the OMM-localized GTPases dynamin-related protein 1 (Drp1) and fission 1 protein (Fis1) (Basu et al., 2017; Smirnova et al., 2001; Yoon et al., 2003). To determine whether loss of Drg1 affects the expression of genes central to these processes, we compared steady-state mRNA levels of a subset of fusion-and fission-associated GTPases (Opa1, Mfn1, Mfn2, Drp1, and Fis1) between wild-type and ΔDrg1 cells. Deletion of Drg1 resulted in increased expression of mRNAs associated with mitochondrial fusion and decreased expression of mRNAs associated with mitochondrial fission. Specifically, Fis1 mRNA levels were reduced by approximately 50%, while Opa1 mRNA levels increased by approximately 1.5-fold relative to wild-type cells (**Figure 6C**). Given the established roles of these factors, the combined increase in fusion-related transcripts and decrease in fission-related transcripts likely contributes to the enlarged, more interconnected mitochondrial network observed in ΔDrg1 cells. Disruption of the balance between mitochondrial fusion and fission, therefore, provides a plausible molecular basis for the abnormal mitochondrial dynamics and morphology observed in the mutant (**Figure 6A**).

ΔDrg1 cells contained fewer and larger mitochondria and exhibited altered network organization, accompanied by changes in the expression of genes involved in mitochondrial fusion and fission. These findings demonstrate that Drg1 loss disrupts mitochondrial architecture and prompted us to determine whether the structural changes are accompanied by impaired mitochondrial function.

### 9. Drg1 is required to maintain mitochondrial membrane potential and ATP production

We next assessed whether the altered mitochondrial network in ΔDrg1 cells was associated with functional impairment by measuring mitochondrial membrane potential and cellular ATP abundance. Mitochondria are essential for ATP production via oxidative phosphorylation. As expected, cellular ATP content was reduced by ∼45% in ΔDrg1 cells compared to WT (**Figure 6D**), suggesting lower ATP production in ΔDrg1. Addition of the WT Drg1 protein back into the ΔDrg1 cells rescued ATP production (**Figure 6D** and **Supplemental Figure S9)**. In contrast, addition of a GTPase-deficient mutant protein, Drg1(S78N), did not restore the amount of ATP to the WT level (**Figure 6D**), suggesting that the catalytic activity of the Drg1 protein is required for the functional rescue. Given the close relationship between ATP synthesis and membrane potential, we tested whether ΔDrg1 cells also exhibited altered membrane potential. Indeed, TMRM staining revealed a ∼30% reduction in ΔDrg1 cells relative to WT (**Figure 6E**). Counterintuitively, the extent of all the above changes is not severe enough to cause measurable differences in cell viability as measured by the MTT assay (**Figure 6F**).

Loss of Drg1 reduced mitochondrial membrane potential and ATP levels without producing a corresponding increase in apoptosis, indicating that the bioenergetic defects are not simply a consequence of widespread cell death. Collectively, these results support a model in which the ribosome-associated translation factor Drg1 regulates the translational engagement and protein output of selected mitochondrial protein–encoding transcripts, thereby contributing to mitochondrial protein homeostasis, architecture, and bioenergetic function.

## Discussion

GTPases are an essential and highly conserved family of proteins that regulate a wide range of fundamental cellular processes (Leipe et al., 2002). Acting as molecular switches, they cycle between GTP-bound and GDP-bound states to control key biological processes, including protein synthesis, cytoskeletal organization, mitochondrial dynamics, vesicle trafficking, and cell signaling. In this study, we demonstrated that Drg, an ancient but poorly understood translational GTPase, is critical for general protein synthesis and for maintaining proper mitochondrial morphology and function. Using multiple complementary approaches, we showed that one of the eukaryotic Drg paralogues, Drg1, is essential for mitochondrial shape, dynamics, membrane potential, and ATP production.

### Drg1 connects ribosome-associated translation to mitochondrial homeostasis

This study identifies mitochondrial homeostasis as an important physiological output of Drg1-dependent translation. Previous biochemical studies showed that mammalian Drg1/Dfrp1 associates with polysomes, whereas structural and functional studies of the yeast ortholog Rbg1–Tma46 established how this conserved complex engages the translating 80S ribosome and promotes efficient translation (Francis et al., 2012; Ishikawa et al., 2009; Zeng et al., 2021). Here, we show that a fraction of the predominantly cytoplasmic human complex associates with the mitochondrial outer membrane, where cytoplasmic ribosomes are also enriched. Mitochondrial-fraction proximity proteomics identified proteins involved in mitochondrial import, translation, oxidative phosphorylation, and proteostasis within the molecular environment of Drg1. Consistent with these associations, loss of Drg1 selectively altered ribosome occupancy on nuclear-encoded mitochondrial transcripts, changed protein output from a mitochondria-targeted reporter, remodeled the mitochondrial network, and reduced mitochondrial membrane potential and ATP production. Together, these findings support a model in which Drg1-dependent cytosolic translation contributes to the production and homeostasis of proteins required for mitochondrial structure and function.

This model provides a mechanistic framework for understanding how a widely distributed cytoplasmic translation factor can produce pronounced effects on a specific organelle. Approximately 99% of the mitochondrial proteome is encoded by the nuclear genome and synthesized by cytoplasmic ribosomes (Rath et al., 2021). Mitochondrial function therefore depends on the coordinated translation, targeting, import, folding, and assembly of proteins produced outside the organelle (Chacinska et al., 2009; Neupert and Herrmann, 2007). Even modest disturbances in these processes may have amplified consequences because oxidative phosphorylation, mitochondrial translation, protein import, and mitochondrial dynamics all depend on the stoichiometric assembly of multiprotein complexes (Chacinska et al., 2009; Neupert and Herrmann, 2007; Rath et al., 2021) Drg1 is therefore positioned to influence mitochondrial function without acting as a conventional mitochondrial-resident protein.

### A mitochondrial-proximal ribosome environment provides a spatial context for a subset of Drg1 function

Our imaging and biochemical fractionation experiments indicate that Drg1 is broadly distributed in the cytoplasm but that a detectable fraction is associated with mitochondria. Proteinase K accessibility further places this population on the cytosolic face of the outer mitochondrial membrane rather than within the organelle. This localization is consistent with a role in cytoplasmic translation and distinguishes Drg1 from mitochondrial matrix GTPases involved directly in mitochondrial ribosome biogenesis or translation.

Cytoplasmic ribosomes were enriched near the mitochondrial surface, providing a plausible spatial context for the mitochondrial-associated Drg1 population. Cytoplasmic ribosomes have been visualized on mitochondrial outer membranes, and proximity-specific ribosome profiling has identified nuclear-encoded mitochondrial mRNAs translated at the mitochondrial surface (Gold et al., 2017; Williams et al., 2014). Recent studies in human cells further demonstrate that a substantial subset of mitochondrial proteins undergoes co-translational targeting or import (Luo et al., 2025; Zhu et al., 2025). Such spatial coupling could reduce the exposure of aggregation-prone precursors in the cytosol and coordinate protein synthesis with mitochondrial import capacity (den Brave et al., 2021; Yamamoto et al., 2009).

Deletion of the Drg1 TGS domain reduced its recovery in mitochondrial fractions, whereas deletion of the HTH domain had a smaller effect. Because the TGS-containing architecture contributes to polysome recruitment of the yeast Rbg1–Tma46 complex (Francis et al., 2012; Zeng et al., 2021), this result is consistent with the possibility that mitochondrial association is mediated, at least in part, through mitochondrial-proximal ribosomes. However, TGS deletion could also alter protein conformation, stability, or interactions independently of ribosome binding. The present findings therefore support, but do not definitively establish, ribosome-dependent recruitment of Drg1 to mitochondria. Experiments that disrupt translating ribosomes and measure the resulting redistribution of Drg1 will be important for testing this model directly.

### The mitochondrial Drg1 environment is linked to protein import and bioenergetic pathways

The APEX2-labeled proteins recovered from mitochondrial fractions provide a molecular view of pathways positioned near mitochondrial-associated Drg1. This dataset included components of the mitochondrial protein-import machinery, respiratory-chain and ATP-synthesis complexes, and mitochondrial ribosomal proteins. The presence of TOMM70A is particularly notable because TOMM70A recognizes subsets of nuclear-encoded mitochondrial precursors, receives precursor–chaperone complexes, and supports delivery to downstream import machinery (Yamamoto et al., 2009; Young et al., 2003). Drg1 proximity to such factors could reflect spatial coupling among cytosolic translation, precursor recognition, and mitochondrial import.

The detection of matrix and inner-membrane proteins does not imply that cytoplasmic Drg1 enters these compartments. APEX2 generates short-lived labeling intermediates and reports molecular proximity rather than direct physical binding (Lam et al., 2015). It may therefore capture nascent mitochondrial proteins before import, newly imported proteins, or mature proteins present in the same biochemical environment. Consequently, the mitochondrial APEX2 dataset should not be interpreted as a set of direct Drg1-binding partners. Furthermore, comparison among biochemical fractions cannot fully distinguish selective Drg1 proximity from differences intrinsic to fraction composition. Nevertheless, when interpreted together with independent imaging, fractionation, and functional measurements, the dataset identifies mitochondrial translation, import, and oxidative phosphorylation as candidate pathways connected to Drg1 activity. Importantly, the APEX2 analysis served as a pathway-level guide for selecting endogenous transcripts for subsequent investigation. This allowed us to move beyond proximity and test whether loss of Drg1 affects the translational state of mRNAs encoding proteins within the implicated mitochondrial pathways.

### Drg1 exerts transcript-selective effects on mitochondrial protein production

Loss of Drg1 did not uniformly alter the total abundance of the mitochondrial protein–encoding mRNAs examined. Instead, it produced transcript-dependent changes in their recovery within the translating ribosome fraction. Ribosome occupancy decreased for several transcripts encoding mitochondrial ribosomal, respiratory-chain, and ATP-synthesis proteins, whereas it increased for others. These results argue against a simple model in which Drg1 globally activates or represses mitochondrial protein synthesis. Rather, Drg1 appears to influence the translational state of many selected mRNA transcripts.

Changes in ribosome occupancy require careful interpretation because ribosome density is a steady-state measurement shaped jointly by initiation, elongation, termination, and ribosome flux (Brar and Weissman, 2015; Ingolia et al., 2009). Reduced occupancy may indicate diminished translation initiation, reduced ribosome stability, accelerated transit, or loss of actively translating ribosomes. Conversely, increased occupancy may reflect increased translational engagement, *slower elongation,* or *ribosome accumulation at difficult sequences.* The results therefore establish altered ribosome behavior but do not, by themselves, quantify completed protein synthesis. Nevertheless, the absence of comparable changes in total mRNA abundance indicates that the effect is not explained solely by altered transcription or steady-state transcript abundance.

### Drg1 supports productive translation through the coding sequence

The reporter experiments reveal a position-dependent defect in protein expression following loss of Drg1. Whereas expression of the reporter positioned toward the 5′ end of the coding sequence was largely preserved in ΔDrg1 cells, the single SOD2-eGFP reporter showed an approximately 5% reduction, and reporter output from the distal 3′ portion of the dual-reporter construct was markedly decreased in a ΔDrg1-specific cell population. This pattern is difficult to explain by a simple defect in transcription, mRNA abundance, or translation initiation, because such mechanisms would be expected to reduce expression of both proximal and distal reporter sequences. Instead, the increasing defect with progression along the coding sequence suggests that Drg1 becomes important after translation has initiated and supports productive expression of distal coding regions.

One possibility is that Drg1 promotes productive translation elongation or ribosome processivity. In this model, ribosomes initiate and translate the 5′ portion of an mRNA relatively normally in ΔDrg1 cells but have an increased probability of slowing, stalling, or failing to complete translation as they progress along the coding sequence. Such a cumulative defect would preferentially reduce protein output from sequences located farther downstream while preserving synthesis of proximal sequences. The stronger phenotype of the 3′ reporter is therefore consistent with a role for Drg1 in promoting successful completion of translation rather than controlling translation initiation.

An alternative, but not mutually exclusive, explanation is translation-coupled mRNA destabilization. Ribosome progression and mRNA stability are intimately linked, and aberrant elongation can engage mRNA surveillance and decay pathways. In ΔDrg1 cells, abnormal ribosome behavior could therefore promote co-translational destabilization or cleavage of the transcript after its 5′ region has already been translated. Such a mechanism would similarly preserve proximal reporter production while disproportionately reducing output from distal coding sequences. In this scenario, altered mRNA stability would not necessarily represent a mechanism independent of translation; rather, it could be a downstream consequence of defective ribosome progression.

We therefore favor a model in which Drg1 supports productive translation through the coding sequence, while the current experiments do not distinguish whether the reduced distal output results directly from impaired ribosome progression, indirectly from translation-coupled mRNA destabilization, or from a combination of the two. Importantly, both mechanisms place the primary consequence of Drg1 loss after translation initiation and predict an increasing probability of unsuccessful protein production as ribosomes progress toward the 3′ end of susceptible transcripts. This model provides a potential framework for understanding the physiological consequences of Drg1 loss at the mitochondrial outer membrane. Localized translation requires not only recruitment of an mRNA and initiation of protein synthesis but also successful elongation and completion of the nascent protein while the translating ribosome is coupled to the mitochondrial surface. Drg1 may therefore protect this later phase of localized protein synthesis, ensuring that initiated translation events yield complete proteins rather than being prematurely interrupted through defective elongation and/or translation-coupled loss of the mRNA template.

### Disruption of Drg1-dependent translation compromises mitochondrial architecture and bioenergetics

Loss of Drg1 produced clear changes in mitochondrial organization, including fewer and larger mitochondria and altered network architecture. These changes indicate remodeling of the mitochondrial network rather than a simple shift in mitochondrial shape. Altered expression of genes involved in mitochondrial fusion and fission may contribute to this phenotype or represent a compensatory response to mitochondrial dysfunction. Mitochondrial morphology is controlled by the opposing fusion and fission machineries and responds dynamically to energetic state, proteotoxic stress, and organelle quality-control pathways (Chen and Chan, 2005; Westermann, 2012); multiple downstream mechanisms may therefore participate.

The structural changes were accompanied by reduced mitochondrial membrane potential and ATP abundance, demonstrating that the phenotype extends beyond morphology to mitochondrial bioenergetic function. Importantly, these defects occurred without a corresponding increase in apoptosis, suggesting that they are not merely secondary consequences of widespread cell death. Instead, mitochondrial dysfunction appears to be an early or direct physiological consequence of Drg1 loss.

The relationship between translation and mitochondrial dysfunction is likely bidirectional. Impaired synthesis or import of mitochondrial proteins can compromise mitochondrial biogenesis, respiration, membrane potential, and cellular proteostasis (Chacinska et al., 2009; den Brave et al., 2021; Neupert and Herrmann, 2007). In turn, reduced cellular energy availability and mitochondrial stress signaling can reprogram cytoplasmic translation and protein homeostasis (Quiros et al., 2016; Suomalainen and Nunnari, 2024). Drg1 loss may therefore initiate a reinforcing cycle in which selective defects in protein production impair mitochondrial function, and mitochondrial dysfunction further challenges the cellular translation and proteostasis systems.

The observed phenotypes are unlikely to arise from failure to produce a single mitochondrial protein. Mitochondrial oxidative phosphorylation, protein import, and organelle dynamics depend on many proteins encoded in both the nuclear and mitochondrial genomes. Modest changes affecting multiple nuclear-encoded components may therefore collectively disrupt the stoichiometry and assembly of mitochondrial complexes. This distributed effect is consistent with Drg1 acting as a translation factor rather than as a mitochondria-specific regulator of one pathway.

### A working model for Drg1-dependent mitochondrial proteostasis

We propose that Drg1-Dfrp1 engages cytoplasmic ribosomes and helps preserve a productive translational state during physiologically challenging elongation events (Saha et al., 2026; Zeng et al., 2021). A subset of these ribosomes operates near the mitochondrial outer membrane, where translation can be coordinated with recognition and import of nuclear-encoded mitochondrial proteins (Gold et al., 2017; Luo et al., 2025; Williams et al., 2014; Zhu et al., 2025). In this setting, Drg1 activity may support synthesis of nascent proteins while limiting prolonged ribosome pausing, premature arrest, or inappropriate engagement of terminal quality-control pathways.

This model does not require Drg1 to recognize mitochondrial transcripts directly. Drg1 could instead respond to a ribosomal state generated during the synthesis of particular proteins. Its apparent transcript selectivity could therefore emerge from features of the coding sequence, nascent chain, translation kinetics, subcellular localization, or the requirement for cotranslational targeting and folding. Determining which of these features recruits or stabilizes Drg1 on individual ribosomes will be an important direction for future work.

Several limitations define the boundaries of the present model. The mitochondrial APEX2 analysis does not distinguish nascent polypeptides from mature neighboring proteins or direct binding partners. Ribosome occupancy does not report elongation rate or completed protein production. Furthermore, TGS-domain dependence is consistent with but does not prove ribosome-mediated mitochondrial recruitment. That said, these limitations concern the precise molecular route connecting Drg1-dependent translation to mitochondria; they do not alter the central observation that Drg1 loss changes the translational state of mitochondrial protein–encoding transcripts and compromises mitochondrial architecture and bioenergetic function.

### Broader implications

The present findings expand the biological significance of the Rbg/Drg family beyond its biochemical activity on the ribosome. They suggest that Drg1-dependent translation contributes to the maintenance of an organelle whose proteome is overwhelmingly produced by cytoplasmic ribosomes. This connection may be especially important in cells with high energetic demands or limited capacity to tolerate disturbances in mitochondrial proteostasis. Recessively inherited loss-of-function Drg1 variants cause a neurodevelopmental disorder characterized by global developmental delay, primary microcephaly, short stature, and craniofacial abnormalities (Westrip et al., 2023). Neurons are particularly dependent on mitochondrial positioning and transport, local protein synthesis, and sustained oxidative metabolism (Holt et al., 2019; Tong et al., 2026). The relationship between Drg1-dependent translation and mitochondrial function identified here therefore provides a potential framework for investigating how altered Drg1 activity contributes to human disease. In summary, by connecting the translational state of cytoplasmic ribosomes to mitochondrial architecture, membrane potential, and ATP production, Drg1 helps coordinate cellular protein synthesis with organellar function.

## Supporting information

Supplemental Table Description

## Acknowledgements

We thank Dr. Boxuan Zhao for valuable guidance on APEX2 labeling. We also thank Sarea Nizami for her help in figure preparation and manuscript proofreading, as well as other members of the Jin laboratory for helpful discussions. We thank Umnia Doha and Glenn Fried at the Institute for Genomic Biology (IGB) for assistance with confocal microscopy and image analysis, and the Roy J. Carver Biotechnology Center at the University of Illinois at Urbana–Champaign for flow cytometry and mass spectrometry support. This research was supported by the National Science Foundation grant to HJ (Award Number: 2408763).

## Materials and Methods

### 1. Cell Culture

HEK293T cells were purchased from The Cancer Center at Illinois, University of Illinois Urbana Champaign (USA). Drg1-KnockOut (ΔDrg1) cells were created using CRISPR-CAS method. Cells were cultured using Dulbecco’s Modified Eagle’s Medium (DMEM) supplemented with 10% FBS, 1 mM pyruvate, and penicillin/streptomycin. Cells were maintained at 37 °C in a humidified atmosphere of 5% CO_2_ as per general cell culture criteria. Cells were transfected using TransIT kit following manufacturer’s protocol.

### 2. Subcellular Fractionation

Cytoplasmic, mitochondrial, and nuclear fractions from HEK293T cells were isolated via a differential centrifugation process (Senichkin et al., 2021). Subcellular fractionation: HEK293T cells were seeded with 1×10^6^ cells per 10 cm dish. At 80% confluency, cells were briefly washed twice with warm PBS, trypsinized and collected in Eppendorf tubes. Cells were washed with ice-cold PBS twice at 500 RCF for 5 minutes at 4 °C. Cells were resuspended in hypotonic buffer (20 mM Tris-HCl (pH 7.5), 10 mM KCl, 2 mM MgCl_2_, 0.5 mM DTT, 0.1% NP-40, Roche Protease Inhibitor Cocktail EDTA Free), passed through a 22-gauge syringe for 10 minutes to prevent clump formation and incubated for 15 minutes on ice. Cells were then centrifuged at 1000 RCF for 5 minutes at 4 °C. The pellet is the nuclear fraction and the cellular debris. The supernatant which is the cytoplasmic fraction is collected and centrifuged at 15000 RCF for 7 minutes at 4° C. The pellet is the mitochondrial fraction, and the supernatant is the cytosolic fraction which is collected separately. The mitochondrial fraction is washed with isotonic buffer (20 mM Tris-HCl (pH 7.5), 150 mM KCl, 2 mM MgCl_2_, 0.5 mM DTT, 0.1% NP-40, Roche Protease Inhibitor Cocktail EDTA Free) twice and centrifuged at 15000 RCF for 5 minutes.

The protocol to sub-fractionate the mitochondrial pellet has been modified slightly (i.e. no NP-40 was added in both hypotonic and isotonic Buffer) from what was previously described (Zhou et al., 2023). The mitochondrial pellet was dissolved in 160 μL of 1X TD buffer (49.9 mM Tris-HCl (pH 7.5), 274.13 mM NaCl, 20.12 mM KCl, ddH2O) as described for mitochondrial outer and inner membrane fractionation. Equal volumes of the mitochondrial fraction were divided into four tubes. To the second, third and fourth tube, we add either 3.3 μL of 20% Triton X-100 or 3.3 μL of Proteinase K (1mg/ml), or both. Incubate the samples at room temperature for an hour with intermittent mixing.

### 3. Fluorescence Microscopy and Raw Image Processing

HEK293T cells were seeded at 0.1×10^6^ cells per well on coverslips placed at the bottom of 6 well plates. For subcellular localization of Drg1 and Dfrp1, cellular mitochondria were labelled with 0.2 μM Mitotracker Red CMXRos (M7512, Molecular Probes) in DMEM for 20 minutes at 37 °C. Cells were then fixed with 4% Paraformaldehyde (PFA) and 0.2% Glutaraldehyde mixture and equal amount equivalent to media in each well was used for 15 mins in standard cell culture condition. The coverslips were washed with PBS and quenched with 50mM Glycine at RT for 20 mins and washed 3 times with 0.1% PBST for 1 min each followed by blocking with 5% Goat Serum at RT for 1 hours. Then coverslips were incubated with primary Anti-Drg1 (Santa Cruz Biotech sc-390620), Anti-DFRP1(Invitrogen Pa5-56637), Anti-RPS6 (Santa Cruz Biotech sc-744559) antibodies overnight in black humidified chamber at 4 degrees. Cells were washed 3 times with 0.1% PBST at RT for 3 mins each and then incubated with Anti-Mouse (Invitrogen A11001) or Anti-Rabbit (Invitrogen A11008) secondary antibodies fused GFP as per primary antibody used followed by 3 times washing with 0.1% PBST, each wash of 3 mins. Slides were prepared using Mounting Dye containing DAPI (Abcam ab104139) to stain nuclei. To measure mitochondrial membrane potential, cells were stained with 0.1 μM TMRM (TMRM, T668, Molecular Probes) and incubated at 37 °C for 30 minutes, washed with PBS, and the coverslips were mounted on slides using mounting solution containing DAPI and imaged instantly. Prepared slides were used for imaging using a LSM900 confocal microscope and Airyscan Imaging (Zeiss). Raw images were Airyscan Processed (for Airyscan images only) and for all images further processing and quantification were done using Fiji. The morphology of mitochondria using 2D-thresholding was analyzed using **mitochondria analyzer** (Fiji, Plug-in). Co-localization study was done using coloc2 (Fiji, Plug-in). For each imaging experiment there were 3 biological replicates. For morphology imaging 4-5 field of view were imaged per replicate. For co-localization study of Drg1 with Mitochondria ∼8 field of view were imaged per replicate and for co-localization study of RPS6 with Mitochondria, 3 field of view were imaged per replicate.

### 4. Cell Viability/Proliferation Assay

MTT assay was conducted using an MTT assay kit, according to the protocol supplied by the manufacturer (Biotium, 30006). Briefly, cells were plated in a 96-well plate at 5000 cells per well and were grown under standard cell culture conditions for 48 hours. Cells were then washed twice with PBS and incubated for 2 hours with 10 μL of MTT in 100 μL of DMEM. After the incubation period, 200μL of DMSO were added to each well and incubated in the dark for another 10 minutes. Absorbance was recorded using BioTek Gen5 plate reader (Agilent) using the manufacturer’s recommended wavelength.6 biological replicates was done.

### 5. APEX Mass Spectrometry and data analysis

HEK293T cells were grown and transfected with plasmid expressing N-terminal V5-APEX-10AA linker-tagged Drg1 gene with Transit (MIR2705, MirusBio) and serum-free media (Gibco) at ∼70-80% confluency. After 24hrs, the transfected cells were prepared for biotinylation using APEX. Cells were treated with Biotin-Phenol (SML2135, Sigma) (final concentration 0.5 mM), containing media. They were incubated at 37 °C for 1 hour, followed by cycloheximide (CHX) (100ug/ml) for 5 minutes. After incubation, hydrogen peroxide (100 mM) was added to the media. For the negative control sample, hydrogen peroxide was omitted. The samples were incubated for 1 minute. The media was aspirated, followed by the addition of quencher (1X Sodium ascorbate (352681000, Thermo), 1X Sodium azide (Thermo, 14314.22) and 1X Trolox (Thermo, 218940010) to the media. Incubate for an additional 1 minute. The cells were washed with the quencher 2 times and then a final PBS wash. Cells were lysed as described in method section titled Subcellular Fractionation. Lysate, mitochondrial, and cytosolic fractions were retained. The cytosolic fraction was further processed in 1M sucrose cushion and spun at 39000 RPM at 4°C for 3 hours and 20 minutes (Beckman Type 45 Ti rotor) to pellet the cytosolic translating ribosomes. The lysate, mitochondria, and polysome samples were incubated with Pierce Streptavidin agarose beads (Thermo, 20353) for one hour and washed with high-salt solution (500 mM). Finally, they were sent for mass spectrometry (UIUC Mass Spec facility) for identification of proteome. The experiment was performed in 3 biological replicates.

To rule out endogenous biotinylation, the lysate of control and experimental samples were used for comparison. Proteins that passed the threshold of log2FC>0 were used for further analysis. To study the polysome interactome of Drg1, lysate vs polysome samples were compared. Proteins with log2FC>1 and p<0.05 were considered enriched in polysome pool. To study the mitochondrial interactome near mitochondria OMM or on mitochondria OMM, the biotinylated enriched proteins from mitochondria and polysome fractions were compared. Similar thresholding values were used to consider mitochondrial enriched hits. The enriched list was subjected to crosscheck against public dataset to highlight nuclear-encoded mitochondrial genes translated on mito-OMM for further downstream experiments. The enriched proteins (gene list) were subjected to GO analysis, and GO-term with FDR<0.05 were selected and plotted as bar plot using R analysis software.

### 6. SDS-PAGE Analysis and Western Blotting

Proteins were resolved by SDS-PAGE and transferred onto nitrocellulose membranes. The appropriate portion of membranes for the molecular weight of each protein were cut, blocked at RT for an 1hour with skimmed milk, followed by incubation with their respective primary antibodies: Anti-DRG1 (Santacruz Biotech sc-390620), Anti-DFRP1 (Invitrogen Pa5-56637), Anti-GAPDH (cytoplasmic marker) (Santacruz Biotech sc-47724), Anti-HSP60 (mitochondrial marker) (Proteintech 15282-1-AP), Anti-TOMM40 (mitochondrial outer membrane marker) (Proteintech 18409-1-AP), Anti-TIMM29 (mitochondrial inner membrane marker) (Proteintech 25652-1-AP), Anti-FLAG (Sigma, F1804), Anti-LSD1 (Nuclear marker) (Santa Cruz Biotech, sc-271720), and Calreticulin (ER marker) (Proteintech,85593-6-RR) at 4 degree overnight. The membrane sections were washed with TBST 3 times, 5 mins each. The aforementioned primary antibodies were all followed by incubation with secondary mouse/rabbit antibody (Thermofisher 31430, 31460) at RT for an hour followed by washing with TBST. The presence of proteins was detected using the ECL detection system (Thermofisher 34577). The membranes were imaged using iBright CL1000 Gel Imager. For Western blot, an equal amount of total proteins were measured using Bradford Assay and loaded into each well on the SDS-PAGE gel.

### 7. Quantitative Real-Time PCR for mito-fusion and fission mRNAs

Total RNA was extracted using the standard TRIzol-based extraction method. 5 μg of total RNA was reverse transcribed using OligodT and Reverse Transcriptase from Sigma (SuperScript III First Strand, Invitrogen) according to the manufacturer’s instructions. qRT-PCR (quantitative Real-Time PCR) was performed by monitoring the real-time increase in fluorescence of POWERUP SYBR Green Master Mix (Applied Bioscience) using a QuantStudio™ Real-Time PCR system (Thermofisher). Samples were analyzed in 3 biological replicates. Each reaction had a final volume of 10 μl and contained 1 × SYBR Green Supermix PCR Master Mix (Applied Bioscience), 0.2 μM forward and reverse primers and 50ng cDNA. PCR primer pairs (IDT) were as follows: Mfn1, 5′-TGTTTTGGTCGCAAACTCTG-3′ and 5′-CTGTCTGCGTACGTCTTCCA-3’; Mfn2, 5′-ATGCATCCCCACTTAAGCAC-3′ and 5′-CCAGAGGGCAGAACTTTGTC-3’; OPA1, 5′-TGTGAGGTCTGCCAGTCTTTA-3′ and 5′-TGTCCTTAATTGGGGTCGTTG-3’; Fis1, 5′-AGGCCGTGCTGAACGAGC-3′ and 5′-GGTAGTTCCCCACGGCCAGG-3’; Drp1, 5′-CACCCGGAGACCTCTCATTC-3′ and 5′-CCCCATTCTTCTGCTTCCAC-3’; GAPDH, 5′-ACATCAAGAAGGTGGTGAAG-3′ and 5′-CTGTTGCTGTAGCCAAATTC-3’.

### 8. qPCR Quantification of Ribosome-bound mRNAs

WT and ΔDrg1 HEK293T cells were grown to 80% confluency and treated with 100 µg/mL CHX for 15 minutes. Cells were harvested by trypsinization, washed twice in ice-cold PBS containing CHX, and resuspended in hypotonic lysis buffer (20 mM Tris-HCl pH 7.5, 5 mM MgCl_2_, 1.5 mM KCl, 0.5% Triton X-100, 1X complete Protease Inhibitor tablet (Roche), 100 U RNase inhibitor, 2 mM DTT). After 20 minutes on ice, cells were lysed by 20 passages through a 22-gauge needle and clarified at 1,000 × g for 5 minutes at 4 °C. KCl was adjusted to 145 mM, and a portion of lysate was layered onto 1 M sucrose cushions and centrifuged at 109,000 × g for 3 hours at 4 °C to isolate ribosome-bound material. Ribosome pellets were resuspended in the polysome buffer (20 mM Tris-HCl pH 7.5, 5 mM MgCl₂, 145 mM KCl, 0.5% Triton X-100, protease inhibitor cocktail, 100 U RNase inhibitor, 2 mM DTT). RNA was extracted from both total lysates and ribosome-associated fractions using TRIzol (Thermo) and reverse-transcribed using SuperScript III (Invitrogen). Quantitative PCR was performed on a QuantStudio instrument (Thermo) using PowerUp SYBR Green Master Mix (Applied Biosystems) and gene-specific primers (GAPDH internal reference). 3 biological replicates were run for each gene. GAPDH, 5′-ACATCAAGAAGGTGGTGAAG-3′ and 5′-CTGTTGCTGTAGCCAAATTC-3’; ATP5D, 5’-ACTCTTCGGTGCAGTTGTTGG-3’ and 5’-GCCTCGATTCGGATCTGGAT-3’; MRPS30, 5’-AGAGCGAGGTCATATCTTTGCC-3’ and 5’-ACCACGCACCCAGTAAAAATG-3’; MRPL43, 5’-TTCTCCACAACGGACTGGGT-3’and 5’-GTCGGGCGAAGTCGATCAC-3’, NDUFB1, 5’-GTCCCTATGGGATTTGTCATTGG-3’ and 5’-CAGTTAGCCGTTCATCACTCTT-3’, ETFA, 5’-CAGAGGAACTGACACCATTGATT-3’ and 5’-AGGTTCTTTCCGAAGGCAGAT-3’, CLPP, 5’-GCCAAGCACACCAAACAGAG-3’ and 5’-TGGACCAGAACCTTGTCTAAGAT-3’, NDUFS4, 5’-TGCTCGCAATAACATGCAGTC-3’ and 5’-GATCAGCCGTTGATGCCCAA-3’, NDUFAF2, 5’-ATGCCTCTGCTCCATACTTTGG-3’ and 5’-TGGCTCTTGCCATCTCGTG-3’, ATP5F1, 5’-GTGAGGACCATGCAAGCC-3’ and 5’-CTGCCAGTTGTTCTCTACTCTGT-3’, UQCRQ, 5’-ATCCGCACGTCTTCACTAAAG-3’ and 5’-TGGATCTCTCGAACTCTTCAGTC-3’, ATP5I, 5’-CAGGTCTCTCCGCTCATCAAG-3’ and 5’-GCCCGAGGTTTTAGGTAATTGT-3’.

### 9. ATP content measurement

Equal numbers of WT and ΔDrg1 HEK293T cells were grown in a 96-well opaque white-walled and bottom plate using DMEM under standard conditions. WT and ΔDrg1 cells were grown for 24 hours. Then a pcDNA plasmid expressing hDrg1 and same backbone expressing hDrg1 with a S78N mutation under CMV promoter were transfected using Trans-IT reagent in separate columns of ΔDrg1 cells. 100ng (**Figure 6D**) and 200ng (**Supplemental Figure S9**) of the mentioned plasmid DNA expressing hDrg1 were used to transfect the same number of cells (∼10,000 cells), in order to see dosage-dependent rescue of the ATP function. 24 hours were given for the plasmids to express in the cells. Then equal volume of CellTiter-Glo (Promega, Catalog Number: G7570) and media were added in each well of the plate. The plate was wrapped in aluminum foil, set for orbital shaking for 5 mins, and then kept aside for 10 mins for the signal to stabilize. Subsequently, luminescence readings representing ATP content were taken using BioTek Gen5 plate reader (Agilent). Eight biological replicates were performed for this analysis.

### 10. eGFP localization, eGFP and mCherry intensity measurement

To study localized translation at mitochondrial OMM, MTS signal sequence of SOD2 was added to eGFP, and 3’UTR of SOD2 gene was added after the stop codon. Another similar construct was made with MTS signal sequence of SOD2 was added to eGFP, after eGFP P2A and mCherry sequence was cloned, and 3’UTR of SOD2 gene was added after the stop codon. These whole constructs were cloned individually into a pcDNA vector. WT cells were grown on cover slip and plasmid was transfected into it, to see expression and localization of eGFP (Valérie Kaltimbacher 2006). Cells were treated with 0.2uM Mitotracker as discussed above. Then the coverslips were mounted on slides using mounting dye containing DAPI. The slides were imaged using LSM900 confocal microscope.

WT and ΔDrg1 HEK293T cells were grown to 80% confluency and transfected with desired construct cloned into a pcDNA plasmid vector using Transit (MIR2705, MirusBio). After 24 hours, cells were washed with PBS, trypsinized, washed 2 times with PBS and resuspended in the PBS solution. The resuspended cells were passed through the flowcytometer; 10000 singlet cells were gated and equal number of cells from both WT and ΔDrg1 cells were identified expressing eGFP. Median eGFP intensity was further calculated, and relative fold change in eGFP compared to WT cells was reported. For the dual fluorescent reporter, 10000 cells expressing GFP was selected, and checked for mCherry expression. The median eGFP and mCherry intensity were calculated and the relative fold change compared to WT cells expressing eGFP and mCherry was reported. The experiment was performed in 3 biological replicates.

### 11. Statistical Analysis

For APEX data, welch t-test was performed with Benjamini-Hochberg correction. For mitochondria morphology data from MitochondriaAnalyzer, qPCR data, ATP content, MTT data, unpaired t-test was performed. For the flow cytometry data, Mann-Whitney U test was performed on the whole data, whereas student t-test was calculated on the median expression. All statistical analyses and plots were done using R software.

## Supplemental Figures and Legends

**Supplemental Figure 1.**
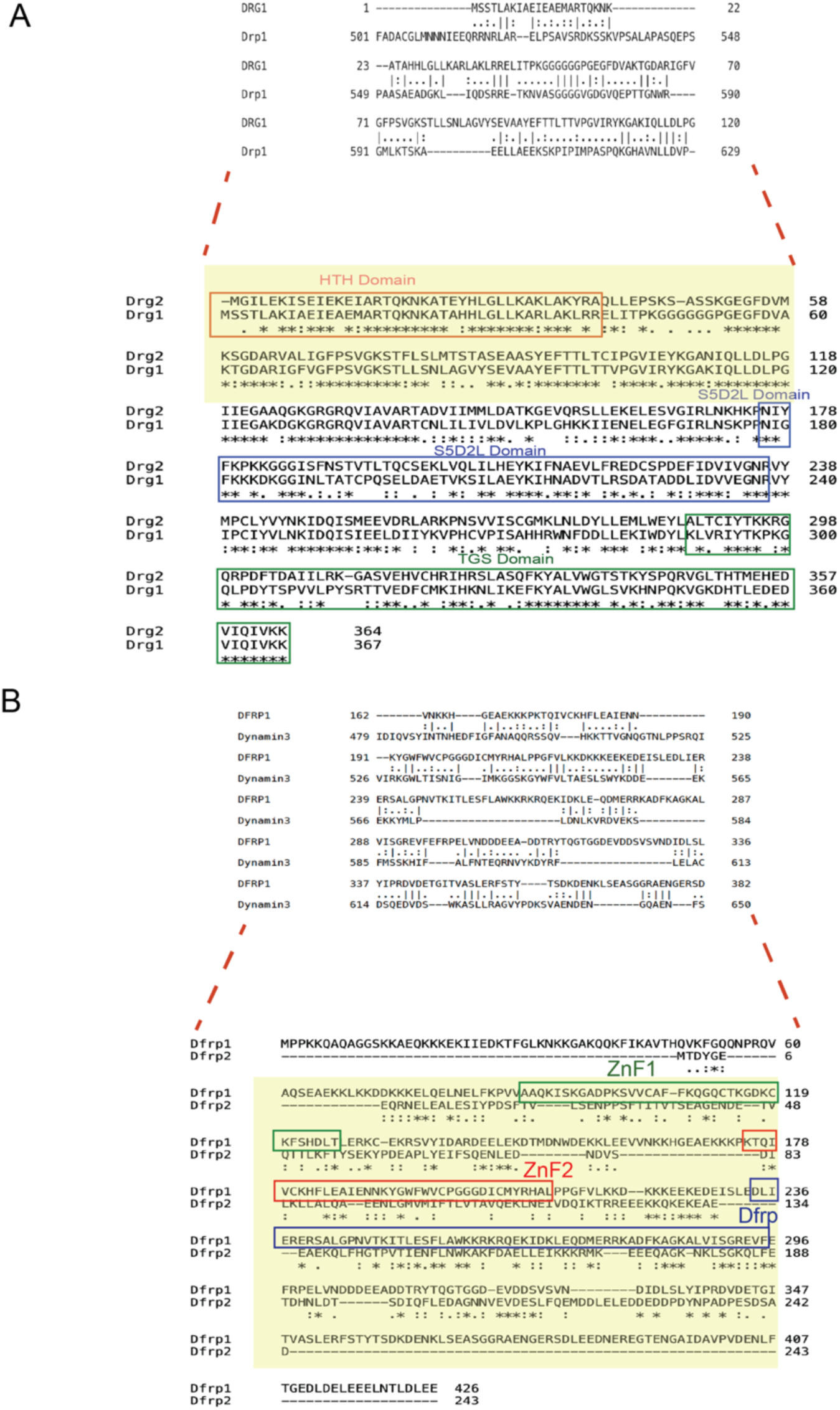
Domain architecture of human Drg and Dfrp proteins. A Human Drg GTPase containing an N-terminal helix-turn-helix (HTH) domain, a central G-domain with an S5D2L insertion, and a C-terminal TGS domain. The TGS domain is essential for binding to Dfrp1 and Dfrp2 and association with the translating ribosome. Drg1 and Drg2 share significant sequence homology with each other. Both HTH and TGS domains are essential for Drg1’s association with the microtubule. Drg1 and Drg2 lack a mitochondrial targeting signal at the N-terminus, suggesting they may not localize into mitochondria. Additionally, Drg1/2 do not share any sequence homology to known lipid-binding domains of mitochondrial outer membrane proteins, as shown in the top panel indicated by the red dashed line. B. Domain architecture of human Dfrp proteins. Dfrp1 has two zinc finger domains Like Drg, Dfrp lacks a mitochondrial targeting signal. Dfrp1 shows no sequence homology to known lipid-binding domains of mitochondrial outer membrane proteins, as shown in the top panel indicated by the red dashed line.

**Supplemental Figure 2.**
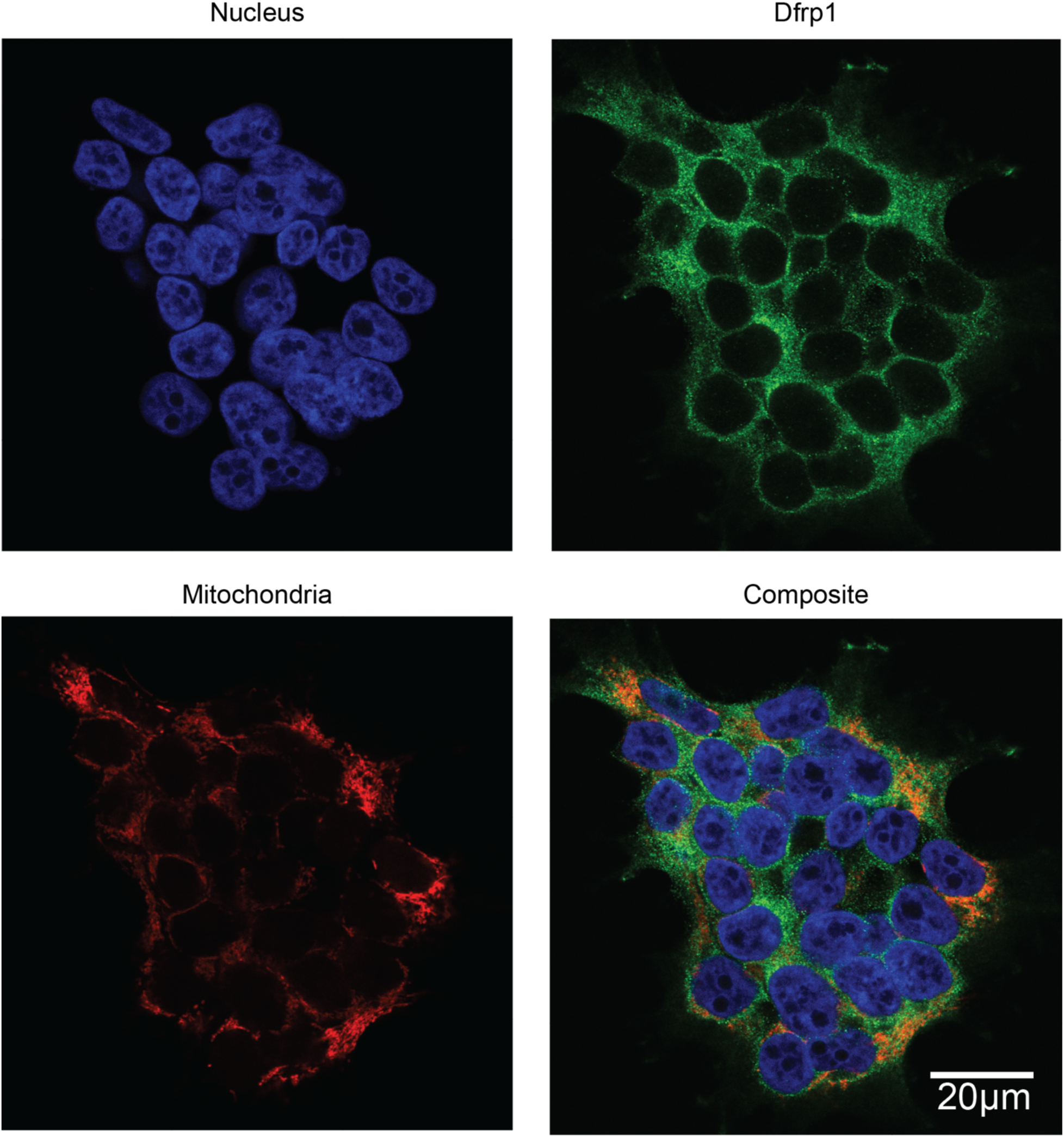
Subcellular localization of Dfrp1. Immunofluorescence confocal imaging of Dfrp1. Nuclei were stained with DAPI (blue), mitochondria with MitoTracker (red), and Dfrp1 with antibodies (green, as indicated). Dfrp1 proteins localize to the cytoplasm.

**Supplemental Figure 3.**
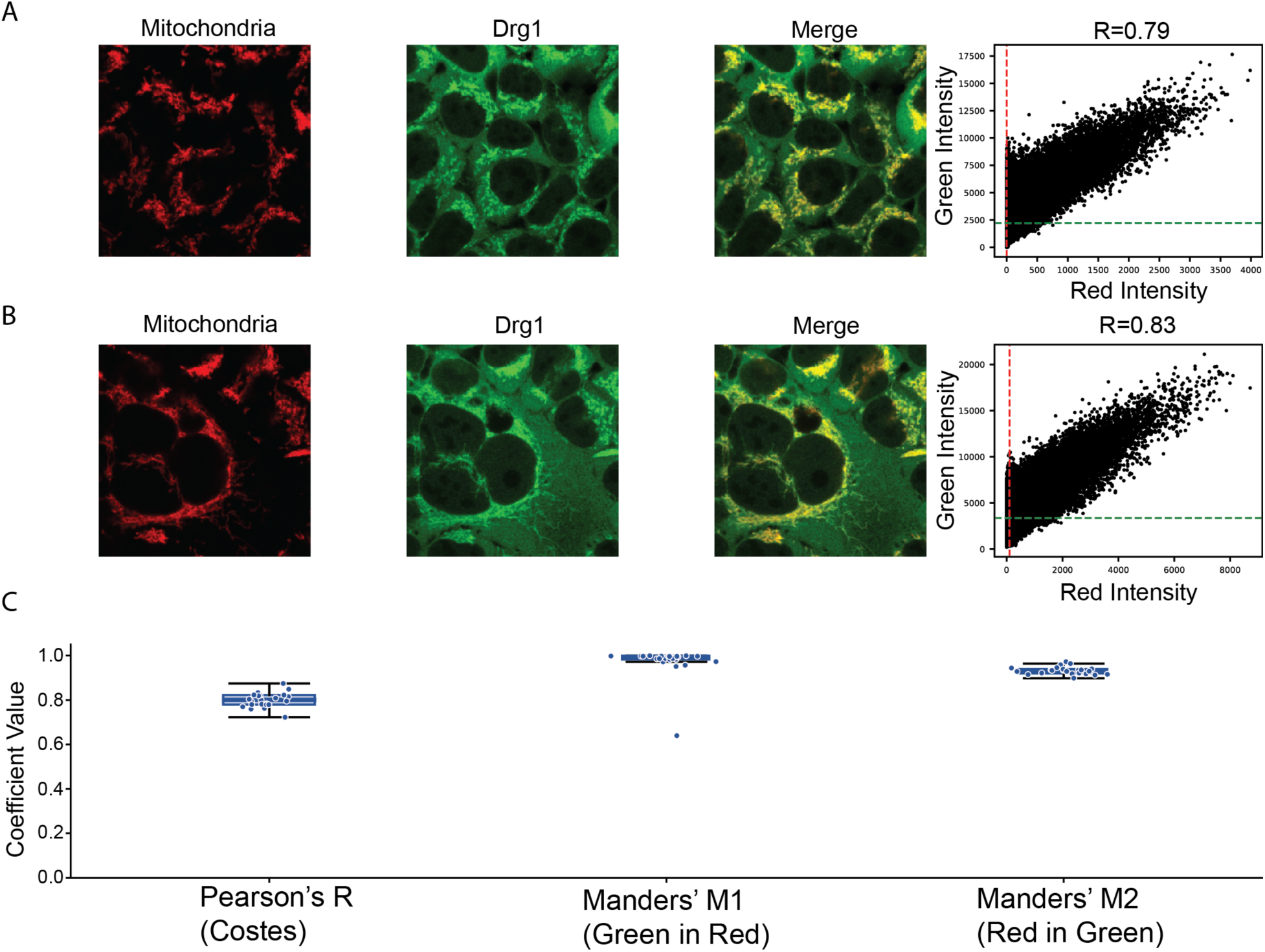
Colocalization of Drg1 with the mitochondrial network. A and B. Representative confocal images of two independent fields showing mitochondria (red), Drg1 immunofluorescence (green), and merge, with pixel-intensity scatter plots and Pearson’s correlation coefficients (R = 0.79 and R = 0.83, respectively). C. Box plots (individual image values overlaid) of Pearson’s correlation coefficient (Costes-thresholded) and Manders’ colocalization coefficients (M1, green-in-red; M2, red-in-green) across all analyzed images.

**Supplemental Figure 4.**
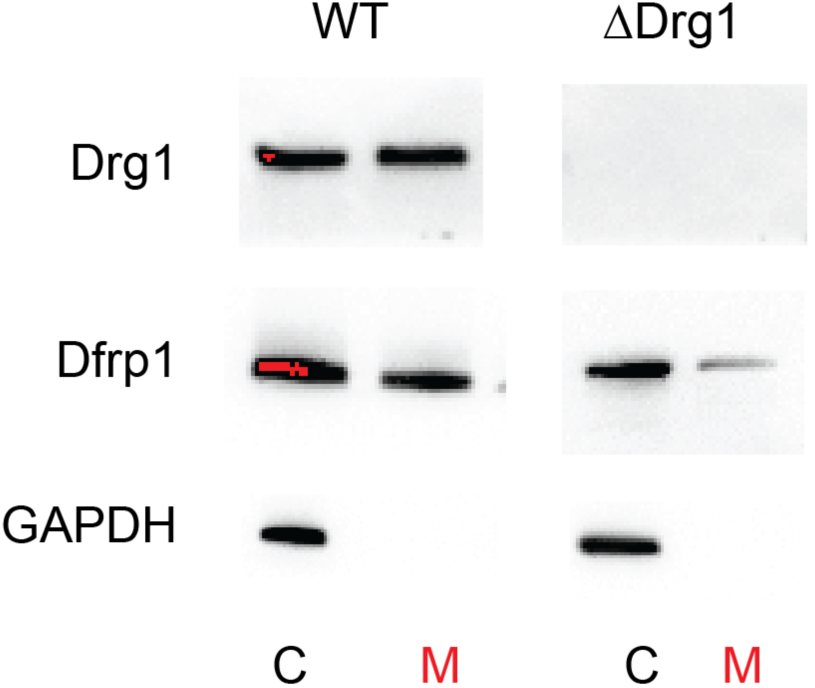
Presence of Drg1 and Dfrp1 in both cytoplasmic and mitochondrial fractions. Western blot analysis of subcellular fractions confirming the presence of Drg1 and Dfrp1 in both cytoplasmic (denoted as “C”) and mitochondrial (denoted as “M”) fractions.

**Supplemental Figure 5.**
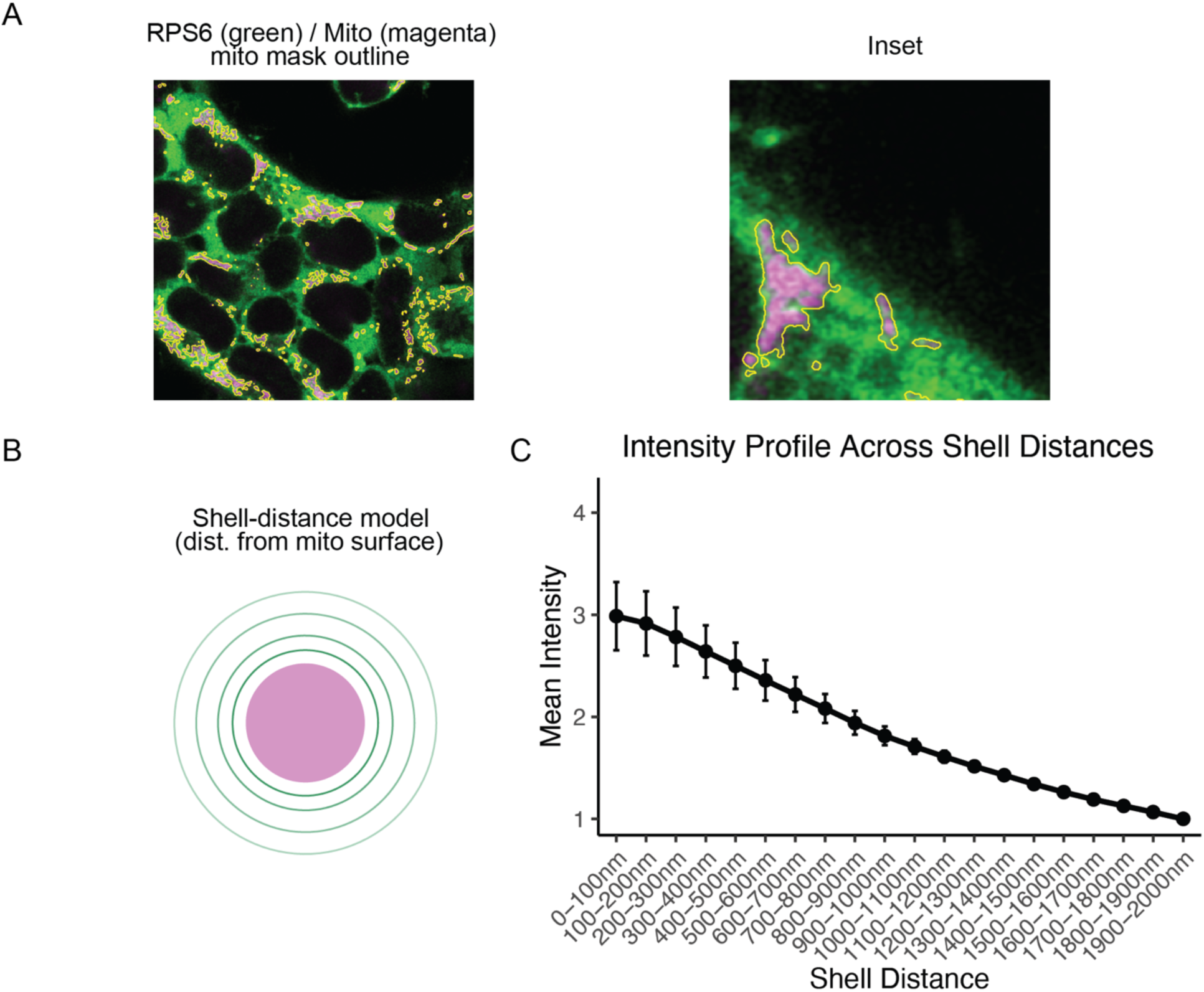
Ribosomal protein RPS6 is enriched in a shell surrounding the mitochondrial surface. A. Representative confocal image of RPS6 (green) and mitochondria (magenta) with the mitochondrial mask outline (yellow) used for shell-distance analysis; right panel shows a magnified inset region. B. Schematic of the shell-distance model used to bin RPS6 intensity by distance from the mitochondrial surface. C. Mean RPS6 intensity as a function of shell distance from the mitochondrial surface (0–2000 nm, 100-nm bins), showing a monotonic decay in RPS6 signal with increasing distance from mitochondria, consistent with ribosome enrichment at the outer mitochondrial membrane.

**Supplemental Figure 6.**
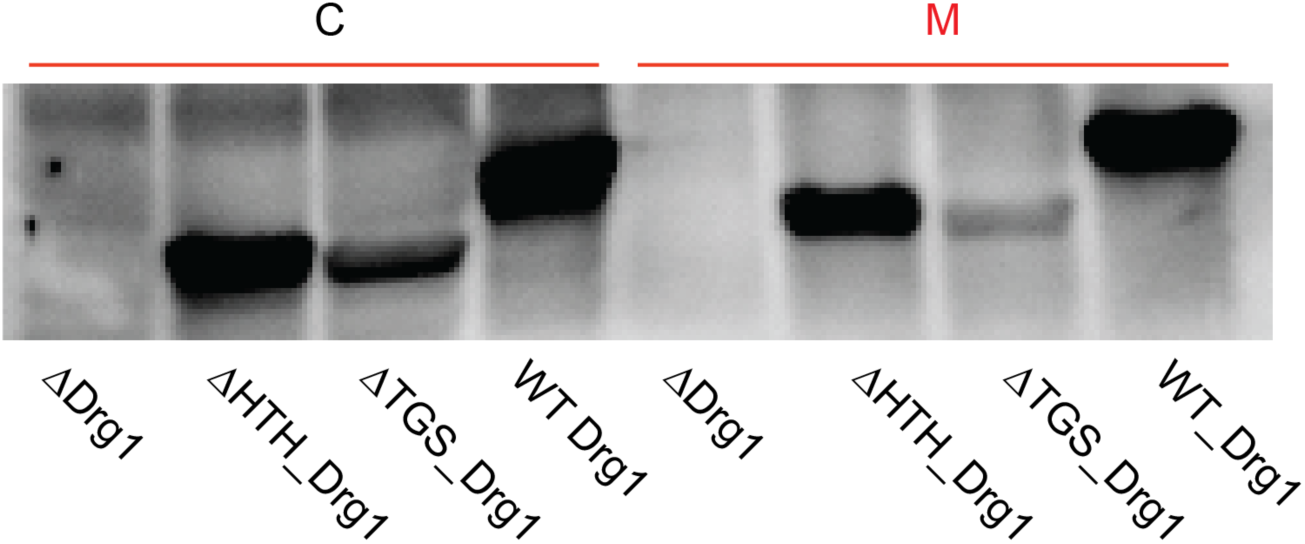
Drg1 domain requirements for mitochondrial localization. ΔDrg1 HEK293T cells expressing Drg1 mutants (ΔHTH or ΔTGS) were imaged as in the panel. The WB was performed by loading equal total amounts of protein for each sample, quantified by Bradford assay. Deletion of the TGS domain abolishes mitochondrial localization, whereas ΔHTH retains localization. These results indicate the TGS domain is critical for Drg1 targeting to mitochondria and/or binding to Dfrp1 and ribosomes.

**Supplemental Figure 7.**
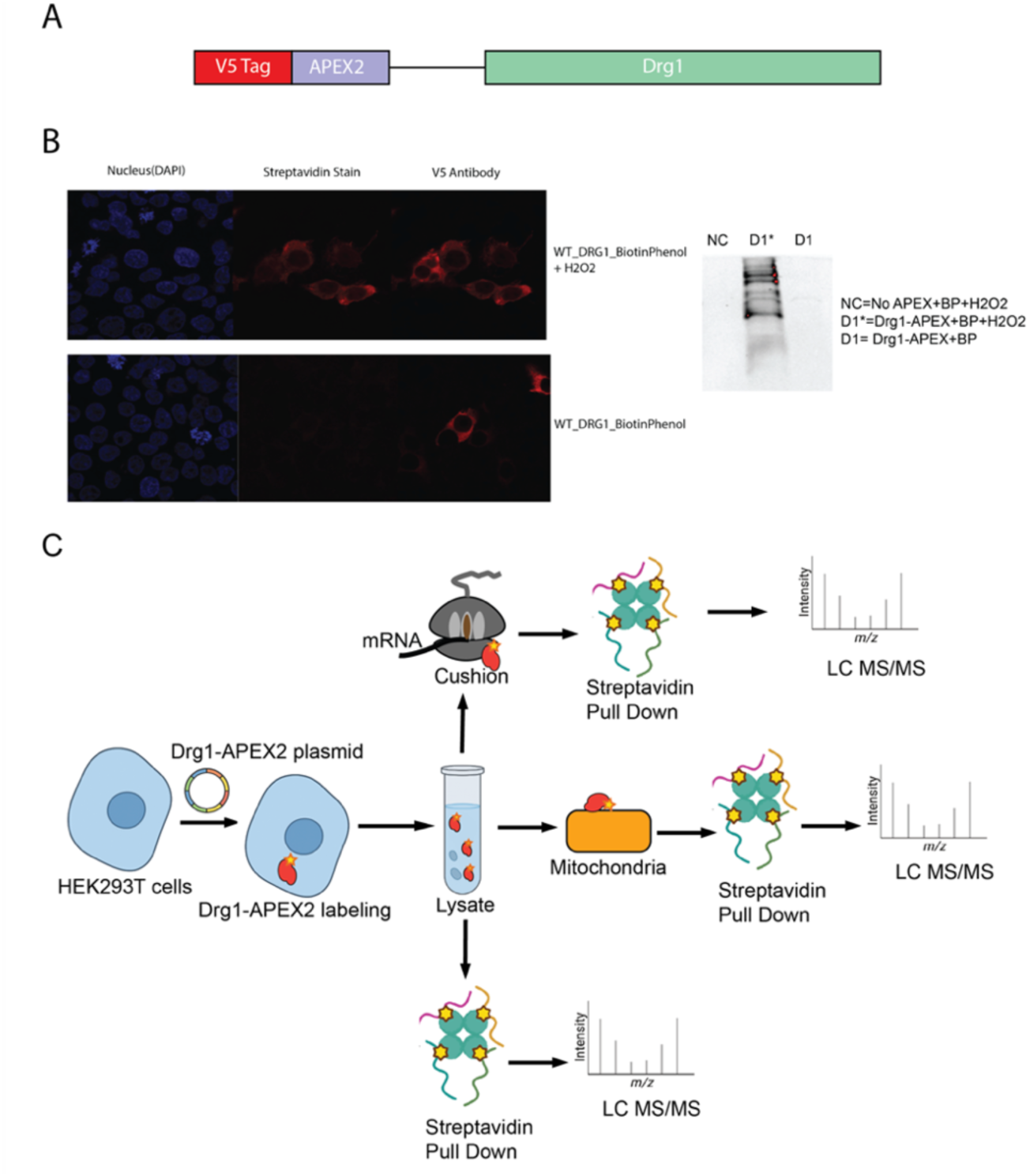
APEX2-Drg1 proximity labeling identifying molecular neighborhood of mammalian Drg1. A. APEX2-Drg1 construct design. V5-APEX2 is attached to the 5’ end of Drg1 with a 10-amino-acid linker in between. This DNA construct is further cloned into a pcDNA plasmid vector under CMV promoter. B. Cytoplasmic localization of APEX2-Drg1. Immunofluorescence imaging of HEK293T cells expressing APEX2–Drg1 constructs shows that the fusion protein localizes predominantly to the cytoplasm, consistent with the localization of endogenous Drg1 (**Figure 1**). Imaging of control and treated samples probed with streptavidin stain confirms that biotinylated proteins are detected only when cells express the APEX2–Drg1 construct and are exposed to both biotin-phenol and H₂O₂, demonstrating the expected conditional labeling. Western blot indicates that the Negative control and no H₂O₂ control showing the biotinylation only work in presence H₂O₂. C. APEX2-Drg1 experimental workflow. Schematic of the Drg1-APEX2 proximity labeling workflow, including APEX2-catalyzed biotinylation, subcellular fractionation, streptavidin pulldown, and LC-MS/MS analysis.

**Supplemental Figure 8.**
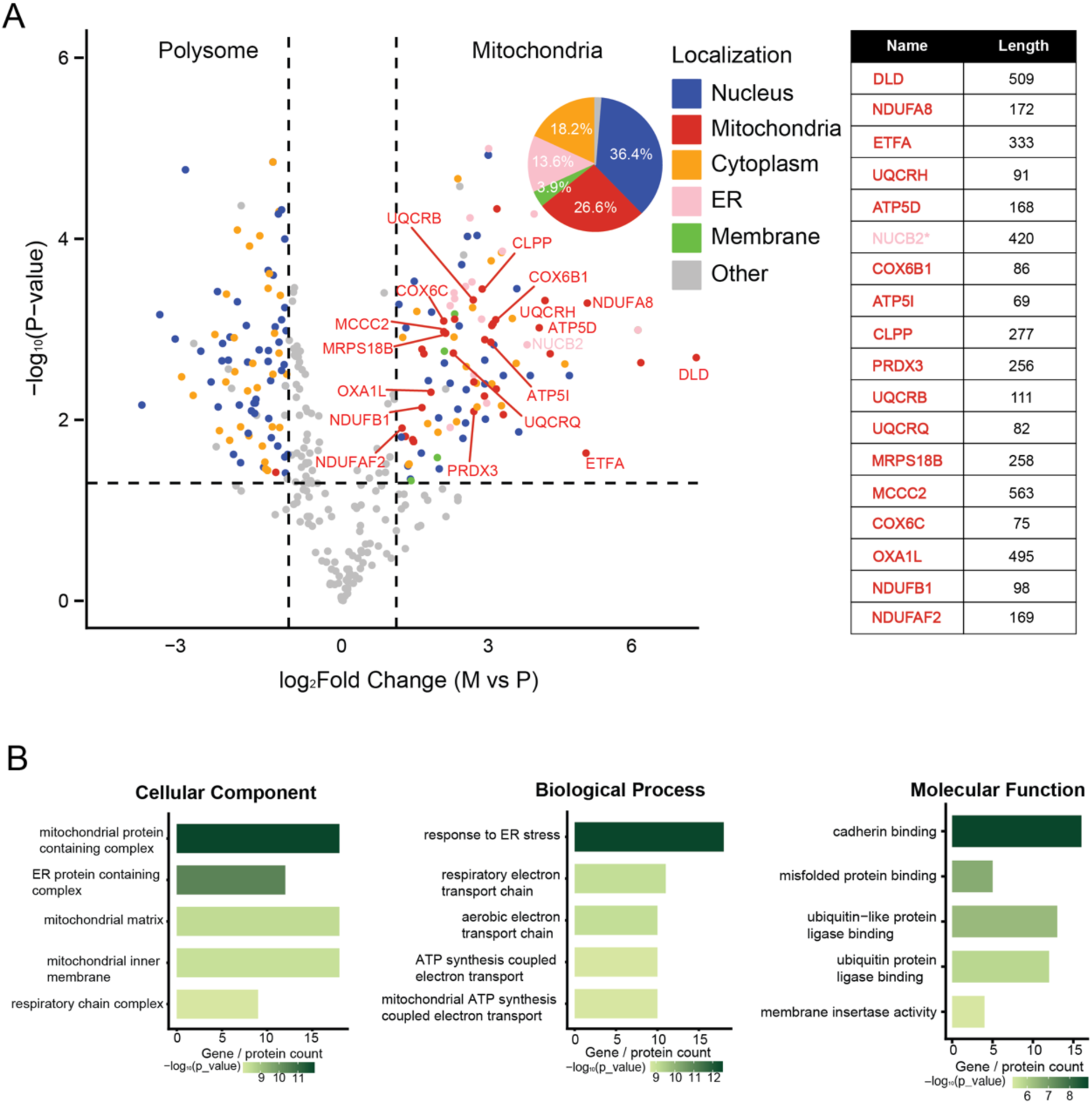
Drg1 interactome at the mitochondrial outer membrane in HEK293T human cells. A Proximity labeling identifies Drg1-associated proteins at mitochondria. HEK293T cells expressing Drg1-APEX2 underwent proximity-dependent biotinylation. Biotinylated proteins were enriched from the mitochondrial fraction and the 80S cytoplasmic ribosome fraction using streptavidin pulldown and identified by LC-MS/MS. The volcano plot displays significantly enriched proteins in the mitochondrial fraction (log_2_ fold change >1, Benjamini-Hochberg adjusted p < 0.05). Proteins known to undergo co-translational mitochondrial import are labeled. Proteins are colored by annotated subcellular localization (Uniprot): cytosol, membrane/ER/Golgi, mitochondrion, nucleus, or other. A subset of identified proteins is shown with their corresponding amino acid (AA) length. All these proteins were reported to undergo co-translational import into mitochondria at the outer mitochondrial membrane, except for NUCB2 (*), which is a protein primarily associated with the endoplasmic reticulum (ER). B. Gene Ontology enrichment analysis of mitochondrial Drg1-interacting proteins, showing terms for cellular compartment, biological process, and molecular function.

**Supplemental Figure 9.**
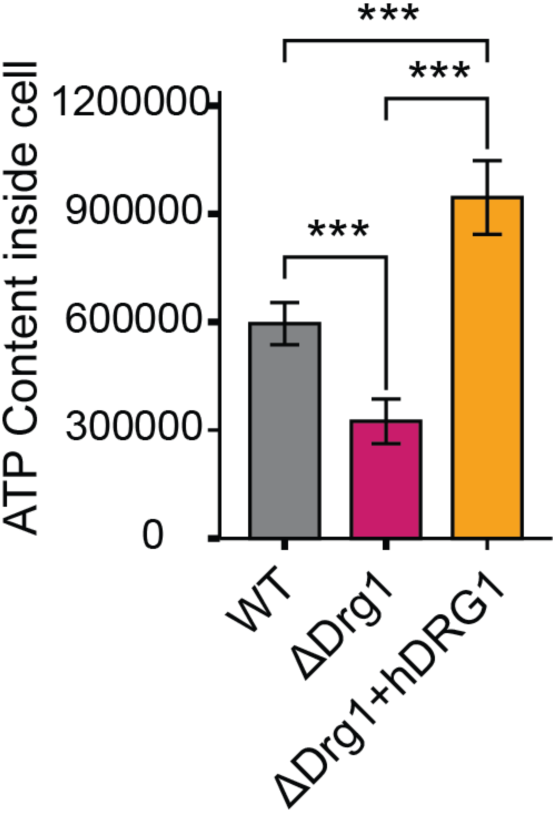
Loss of Drg1 reduces cellular ATP levels. Relative cellular ATP levels in WT, ΔDrg1 cells, as well as ΔDrg1 cells with WT Drg1 expressed. Loss of Drg1 reduces ATP levels, and the phenotype was rescued upon expressing WT Drg1 back into the ΔDrg1 cells. In this experiment, 200ng of plasmid DNA containing WT human Drg1 gene was transfected into ∼10,000 cells.

