## Supplemental Table Description for "Ribosome-Associated GTPase Drg1 Links Cytosolic Translation to Mitochondrial Homeostasis"

**Supplemental Table 1. Proteomics analysis of Drg1-interacting proteins enriched in mitochondrial versus polysome fractions.**

Data presents the enriched dataset of 154 proteins identified in the Drg1 interactome, comparing mitochondrial and polysomal fractions, with $\log_2$Fold-Change>1 and p-value<0.05. The enriched list of genes highlighted in yellow has known mitochondrial localization and function according to the MitoCarta 3.0 database [1]. Among them, 23 genes in **bold** are reported to undergo co-translational import into mitochondria [2,3].

Columns report gene name, UniProt ID, protein names, subcellular locations, function, molecular weight (kDa), amino acid sequence length, label-free quantification (LFQ) intensities for proteins detected in both polysomal and Mitochondrial fractions in three biological replicates, with $\text{Log}_2$Fold-Change, p-value
